# Anaerobic Benzene Biodegradation Linked to Growth of Highly Specific Bacterial Clades

**DOI:** 10.1101/2021.01.23.427911

**Authors:** Courtney R. A. Toth, Fei Luo, Nancy Bawa, Jennifer Webb, Shen Guo, Sandra Dworatzek, Elizabeth A. Edwards

## Abstract

Reliance on bioremediation to remove benzene from anoxic environments has proven risky for decades but for unknown reasons. Years of research have revealed a strong link between anaerobic benzene biodegradation and the enrichment of highly specific microbes, namely *Thermincola* in the family Peptococcaceae and the deltaproteobacterial Candidate Sva0485 clade. Using aquifer material from Canadian Forces Base Borden, we compared five bioremediation approaches in batch microcosms. Under conditions simulating natural attenuation or sulfate biostimulation, benzene was not degraded after 1-2 years of incubation and no enrichment of known benzene-degrading microbes occurred. In contrast, nitrate-amended microcosms reported benzene biodegradation coincident with significant growth of *Thermincola* spp., along with a functional gene presumed to catalyze anaerobic benzene carboxylation (*abcA*). Inoculation with 2.5% of a methanogenic benzene-degrading consortium containing Sva0485 (*Deltaproteobacteria* ORM2) resulted in benzene biodegradation in the presence of sulfate or under methanogenic conditions. The presence of other hydrocarbon co-contaminants decreased rates of benzene degradation by a factor of 2-4. Tracking the abundance of the *abcA* gene and 16S rRNA genes specific for benzene-degrading *Thermincola* and Sva0485 is recommended to monitor benzene bioremediation in anoxic groundwater systems to further uncover growth rate limiting conditions for these two intriguing phylotypes.

**SYNOPSIS:** Anaerobic benzene biodegradation was accelerated by biostimulation with nitrate or by bioaugmentation under methanogenic or sulfate-reducing conditions.

## 1 Introduction

Benzene, toluene, ethylbenzene, and xylenes (BTEX) are widespread contaminants owing to petroleum spills and releases at industrial facilities, oil refineries, underground storage tanks, or from pipelines and mining operations. Compared to other hydrocarbons in petroleum, BTEX are relatively water soluble and can be transported far from the original source leading to extensive contamination. Cleanup is generally dictated by feasibility and economic viability. For example, shallow spills can be more easily excavated and/or aerated, and usually rely on aerobic (bio)remediation processes for complete contaminant destruction.^1^ Low permeability groundwater zones, deep aquifers and sediments, and sites below existing infrastructures often require *in situ* technologies and thus rely more significantly on anaerobic biodegradation.^2^ While the latter has been an effective remediation approach for toluene and xylenes,^3-7^ anaerobic biodegradation of benzene has been largely unreliable. Only a handful of known field reports have ever demonstrated convincing evidence of anaerobic benzene bioremediation ^3,8,9^ despite numerous isotopic analyses supporting that anaerobic benzene degradation does occur *in situ*.^10-13^ This poses a challenge for site managers, as benzene is often the driver for remediation efforts due to its confirmed carcinogenicity and lowest allowable concentrations in groundwater (≤ 5 µg/L in Canada and the US).^14^

Anaerobic benzene-degrading microbes identified to-date are mostly uncultured strains and belong to only a few clades primarily within the Deltaproteobacteria and the Firmicutes (Figure 1). Their roles in anaerobic benzene oxidation have been gleaned through stable isotope probing ^15-17^, using metagenomic and related approaches ^18-22^, enrichment and clone sequencing ^23-25^, and growth studies in the case for isolated strains of *Geobacter* ^26^. Phylotypes associated with benzene cluster with other anaerobic polycyclic aromatic hydrocarbon (PAH) degraders and based on electron acceptor (Figure 1). Here, we will briefly overview the two most consistently-documented benzene-degrading clades: the Candidate Sva0485 order of *Deltaproteobacteria* and the genus *Thermincola* within the *Peptococacceae*.

**Figure 1.**
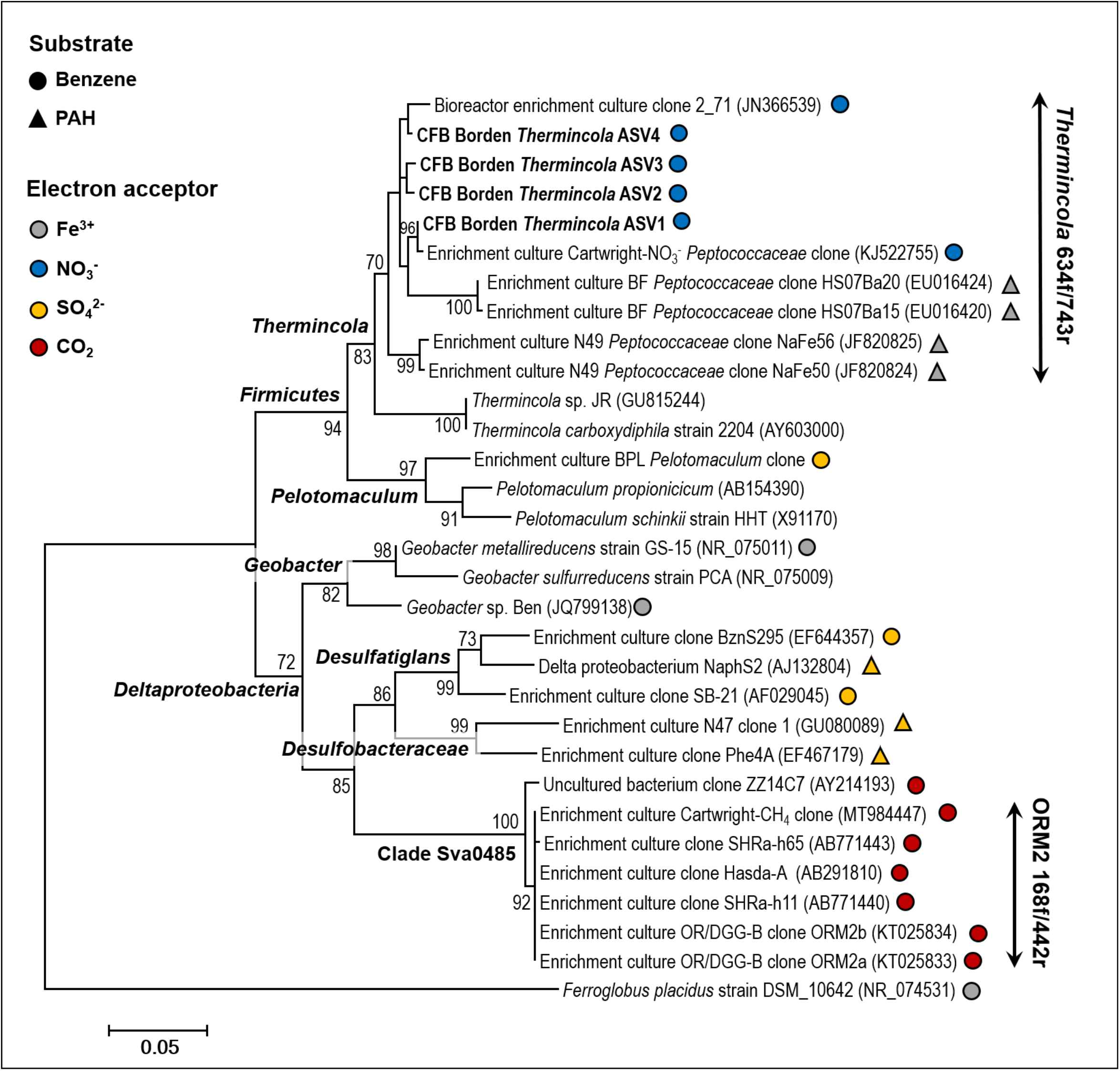
Maximum likelihood consensus tree showing the affiliation of near-complete 16S rRNA genes (1231 bp) belonging to anaerobic benzene and polycyclic aromatic hydrocarbon (PAH)-degrading microorganisms, and select reference strains. A description of how the final consensus tree was constructed is provided in Supporting Information (Text S1). Additionally, the specificity of the *Thermincola* and ORM2 qPCR primer pairs used in this study is illustrated.

### 1.1 Deltaproteobacterial Candidate Sva0485

In studies of methanogenic benzene-degrading enrichment cultures dating back to 1995, our laboratory postulated that a 16S rRNA gene sequence clone of a deltaproteobacterium referred to as ORM2 belonged to the benzene degrader in the OR consortium derived from an Oklahoma oil refinery.^25, 27^ Further proof came in subsequent studies, where Da Silva and Alvarez^28^ inoculated laboratory aquifer columns with the OR consortium, and used a combination of denaturing gradient gel electrophoresis and 16S rRNA gene-based quantitative PCR (qPCR) tests to confirm significant proliferation of *Deltaproteobacteria* ORM2. Growth of ORM2 was always coincident with the establishment of anaerobic benzene biodegradation activity. Similarly, the authors observed a rapid decrease in *Deltaproteobacteria* ORM2 copies shortly after benzene was depleted.^28^ Later, metagenomic surveys found the existence of two closely related strains of ORM2 (ORM2a and ORM2b) in the OR consortium, both of which were hypothesized to degrade benzene fermentatively with methanogens and possibly coupled to sulfate reduction.^20, 29^ Luo et al.^20^ confirmed the relationship between 16S rRNA gene copy abundance of both *Deltaproteobacteria* ORM2 strains and benzene biodegradation activity in multiple OR subcultures. Refined 16S rRNA databases now place ORM2 within the recently proposed Candidate Sva0485 clade (Figure 1). This clade also includes several other benzene-associated fermenters from different geographical origins. For example, an ORM2-like strain exists in a second methanogenic benzene-degrading consortium maintained in our laboratory (Cartwright-CH_4_), that was enriched from a decommissioned gasoline station in Toronto, Ontario, Canada.^20, 27^ Research groups from Japan have also identified nearly identical 16S rRNA sequences to ORM2 (designated Hasda-A and SHRah65) in parallel long term benzene enrichment culture studies.^17, 30^ Most recently, Qiao et al.^8^ demonstrated convincing field and microcosm evidence that anaerobic benzene bioremediation at a contaminated industrial site in China was attributed to enrichment of intrinsic Sva0485 Deltaproteobacteria. No member of the Candidate Sva0485 has yet to be isolated, and the mechanism(s) for benzene activation by these microorganisms is unknown. Interestingly, no substrate other than benzene has been found to support growth of *Deltaproteobacteria* ORM2/Candidate Sva0485 organisms in mixed cultures, including benzoate, phenol, or toluene.^31 29^This is consistent with the annotations of a complete metagenome-assembled genome of deltaproteobacterium ORM2a available at JGI/IMG (taxon ID 2795385393).

### 1.2 Thermincola

Benzene-degrading *Thermincola* species have been enriched under iron-reducing and nitrate-reducing conditions, and have been retrieved from various materials in Canada, Poland, and the Netherlands.^15, 16, 25^ Ulrich and Edwards^25^ first identified identical *Thermincola* 16S rRNA gene sequence clones in nitrate-reducing benzene-degrading cultures from the aforementioned gasoline station in Toronto (Cartwright-NO_3_^-^, see Figure 1) and from an uncontaminated swamp near Perth, Ontario, Canada. The role of *Thermincola* in benzene degradation was later supported by Kunapuli et al.^15^ and van der Zaan et al.,^16^ who each identified a handful of isotopically-labelled *Thermincola* 16S rRNA gene sequence clones after feeding mixed iron-reducing and nitrate-reducing cultures, respectively, with ^13^C_6_-benzene. Using the enrichment culture BF (Figure 1), Abu Laban et al.^22^ employed metagenomics and differential comparative proteomics analyses to identify a functional gene (*abcA*) encoded by *Thermincola* predicted to catalyze the direct carboxylation of benzene. A metatranscriptomic analysis of the Cartwright-NO_3_^-^ culture (Luo et al, 2014) also supported a role for carboxylation in benzene activation. Quantitative PCR assays targeting *abcA* have since been developed by our lab and at least two other research groups,^32-35^. Notably, the *abcA* gene has only ever been identified in benzene-degrading *Thermincola* species, suggesting this mechanism of benzene activation may be clade-specific. From metagenome sequencing of the Cartwright-NO_3_^-^ culture from our laboratory, a draft genome of *Thermincola* in 27 contigs (JGI/IMG taxon ID 2835707023) has recently been obtained. Most likely, *Thermincola* initiates benzene biodegradation in a fermentative process and downstream metabolites are further transformed by several other groups of microbes, primarily denitrifying microbes such as *Aromatoleum* and other betaproteobacteria. Attempts to grow the benzene-degrading consortium on any substrate other than benzene have so far only managed to enrich downstream taxa.^31^

### 1.3 Implications and Research Objectives

If specific microbial clades are responsible for anaerobic benzene attenuation, then strategies to increase the abundance of these organisms *in situ* must be identified to enhance bioremediation. Developing specific biomarkers that can reliably monitor the abundance known benzene degraders is also imperative. Because all predicted fermentative benzene-degrading organisms belong to Candidate Sva0485 (Figure 1), and because the only catalytic gene available predicted to encode for anaerobic benzene carboxylase (*abcA*) has been exclusively identified in benzene-degrading *Thermincola*, we hypothesized that qPCR methods to track the presence and abundance of these two organisms and the *abcA* gene might provide the specificity needed to infer benzene biodegradation at contaminated sites.

To this end, we compared five bioremediation approaches (natural attenuation, sulfate biostimulation, nitrate biostimulation, bioaugmentation, and a combination of bioaugmentation and sulfate) in a series of microcosms constructed with sediments from a petroleum-contaminated aquifer in Ontario, Canada. This site was selected because previous isotopic and qPCR evidence suggested intrinsic anaerobic benzene degradation coincident with increases in *abcA* gene copies.^32^ Our primary objective was to identify strategies that could reliably enrich active anaerobic benzene degraders, either those naturally present in site sediments or artificially inoculated from a defined methanogenic benzene degrading culture. Since benzene is rarely the sole pollutant at contaminated sites, a second objective was to explore the impact of related co-contaminants (toluene, ethylbenzene, *o*-xylene, *m*-xylene and naphthalene) on anaerobic benzene degradation. Our results show that intrinsic benzene-degrading *Thermincola* could be enriched only in the presence of nitrate, and that bioaugmentation (with the OR consortium containing *Deltaproteobacterium* ORM2) stimulated benzene removal under methanogenic and sulfate-reducing conditions. The presence of co-contaminants (TEX and naphthalene) led to longer removal timeframes in these microcosm studies.

## 2 MATERIALS AND METHODS

### 2.1 Description of the Methanogenic Benzene-Degrading Culture DGG-B

The oil refinery (OR) consortium was established in 1995 from hydrocarbon-contaminated soil and groundwater samples collected at an oil refinery site in Oklahoma, USA.^25, 27^ For approximately 20 years, subcultures of the parent microcosms were maintained as described elsewhere^20, 25^ and repeatedly fed benzene (130 – 1,100 µM) as their sole carbon and energy source. A schematic of the subculturing history of the OR consortium is available in Luo et al.^20^ In 2016, one active subculture (OR-b1Ar) was transported to SiREM labs (Guelph, ON) for scale-up to produce commercial volumes (>100L), which we now refer to as the DGG-B lineage, named in honour of anaerobic hydrocarbon biodegradation pioneer, Dunja Grbić-Galić. In DGG-B culture vessels, rates of benzene degradation have averaged 1.4 – 25 µM/day.

### 2.2 Sample Location and Collection

Groundwater aquifer sediments were collected in September 2016 from a shallow, petroleum hydrocarbon-contaminated aquifer at the Canadian Forces Base (CFB) Borden in Ontario, Canada. The site is managed in part by the University of Waterloo, where it is used for field experimentation of groundwater remediation technologies.^32, 36^ The water table is located 1.0 meter below surface (mbs), varying seasonally, and contains background concentrations of total iron (25 – 30 mg/L) and sulfate (10 to 30 mg/L) supplied from a nearby historical leachate plume.^37, 38^ At the time of sampling, groundwater concentrations of benzene, toluene and xylenes ranged between 200 – 960 µg/L. Sediment cores were retrieved from a depth of 1.5 – 3.0 mbs, encased in plastic tubing, and capped to minimize sediment exposure to oxygen. The extracted sediments were homogenous in appearance and composed primarily of fine to medium grain sand. All samples were immediately shipped to SiREM (Guelph, ON) and stored at 4 °C until use.

### 2.3 Experimental Setup

In a disposable glove bag (Atmosbag, Sigma-Aldrich), homogenized CFB Borden sediments were distributed (60 g) into 250-mL glass screw-cap bottles, followed by 150 mL of anaerobic artificial groundwater.^39^ Resazurin was also added to one bottle per experimental treatment (described below) as a redox indictor. Bottles were sealed with Teflon Mininert^®^ valve screw caps and stored in an anaerobic glove box (headspace supplied with 10% H_2_, 10% CO_2_, 80% N_2_) for two weeks prior to any further amendments to ensure anaerobicity.

On Day 0 of the experiment, prepared microcosms were amended with electron acceptors and/or electron donors, as detailed in Table S1. Three to four bottles were established per experimental treatment, for a total of 38 microcosms. Microcosms were named numerically and include the abbreviation BOR in reference to CFB Borden. On Day 7, select microcosms were bioaugmented with 3.75 mL (2.5% *v/v*) of DGG-B culture. Microcosms were incubated for up to 645 days and were sampled approximately every four weeks as outlined below. Specific bottles were re-amended with respective electron donor(s) and/or electron acceptor when depleted.

### 2.4 Analytical Methods

Methane, BTEX, and naphthalene dissolved in the aqueous phase of the microcosms were routinely measured using an Agilent 7890 gas chromatograph (GC) equipped with an Agilent G1888 headspace autosampler, described in detail in Supporting Information (SI). Anion (nitrate, nitrite, sulfate, acetate equivalents, chloride, and phosphate) analysis was performed on a Thermo-Fisher ICS-2100 ion chromatograph (IC) equipped with a Thermo-Fisher AS-DV autosampler and an AS18 column (SI Text S2). Iron reduction was not monitored in this study. Raw data are reported in Table S2.

### 2.5 DNA extractions and analyses

Microcosms were sampled regularly for subsequent genomic DNA (gDNA) extraction and analysis to monitor growth of known anaerobic benzene degraders and microbial community composition. Briefly, solid and liquid slurries (1 mL) from well-shaken microcosms were centrifuged for 5 min at 13,000 × *g* and their supernatants discarded. The resulting pellets were frozen (−80 °C) and genomic DNA (gDNA) extracted using the KingFisher^™^ Duo Prime Purification System (Thermo Scientific).

All gDNA samples were assayed by qPCR using universal 16S rRNA gene primer sets for Bacteria and Archaea, as well as specific primers for Sva0485 clade Deltaproteobacteria (including ORM2) and benzene-degrading *Thermincola* (Table S3). We also tracked the predicted anaerobic benzene-carboxylase gene *abcA* (Table S3). Reaction efficiencies for all primer sets ranged between 87 and 105% and calibration curve R^2^ values were > 0.995 (Figure S1). The specificity of the ORM2 and *Thermincola* primer pairs against sequences of known/predicted benzene and PAH-degrading bacteria is indicated in Figure 1. qPCR reaction and thermocycling conditions are available in SI (Text S3). Quantification limits for qPCR were approximately 10^3^ copies per mL of microcosm slurry.

The relative abundance of other microbes in each sample was determined using 16S rRNA gene amplicon sequencing using primers targeting the V6-V8 region. Details are provided in SI (Text S4). Raw sequence reads were deposited to the National Center for Biotechnology Information Short Read Archive (SRA) under BioProject PRJNA661350. No 16S rRNA gene amplicons were recovered from sterile poisoned control bottles at any timepoint.

## 3 RESULTS AND DISCUSSION

### 3.1 Anaerobic benzene biotransformation and microbial abundances by treatment

Thirty-eight (38) anoxic microcosms containing CFB Borden (BOR) sediments were established. At the start of the experiment (Day 0), all microcosms initially contained small amounts of nitrate (0.8 ± 2.0 µmol/bottle), sulfate (23 ± 2.0 µmol/bottle) and methane (8.0 ± 0.2 µmol/bottle) from site sediments (Table S2). Deltaproteobacterial Candidate Sva0485 16S rRNA genes (< 10^3^ copies/mL) and *Thermincola* 16S rRNA genes (≤ 10^4^ copies/mL) detected by qPCR in the gDNA extracted on Day 0 were at or just above detection limits, suggesting that very low numbers of intrinsic anaerobic benzene degraders in BOR sediments, in agreement with previous field reports.^32^

Following addition of hydrocarbon electron donors, electron acceptors, and inoculation with DGG-B culture as per Table S1, concentrations were monitored over time. The main trends observed for each treatment are also summarized in Table S1; all experimental raw data are provided in Table S2. Benzene was not depleted in mercuric chloride-poisoned sterile controls (Figure S2a). Benzene was also not degraded in untreated (BOR05-07; Figures 2a) or 2 mM sulfate-amended microcosms (BOR08-10; Figure 2b), despite active methanogenesis or sulfate reduction in these bottles. Based on qPCR analyses, intrinsic Sva0485, *Thermincola* and *abcA* gene copies did not increase above quantifiable limits in either set of treatments (Figures 2a and 2b; bottom panel). Even after 645 days, no convincing evidence of benzene depletion was observed (Figures S2b andc). Given the overall lack of benzene degradation, no additional molecular timepoints were collected in these bottles after Day 109.

**Figure 2.**
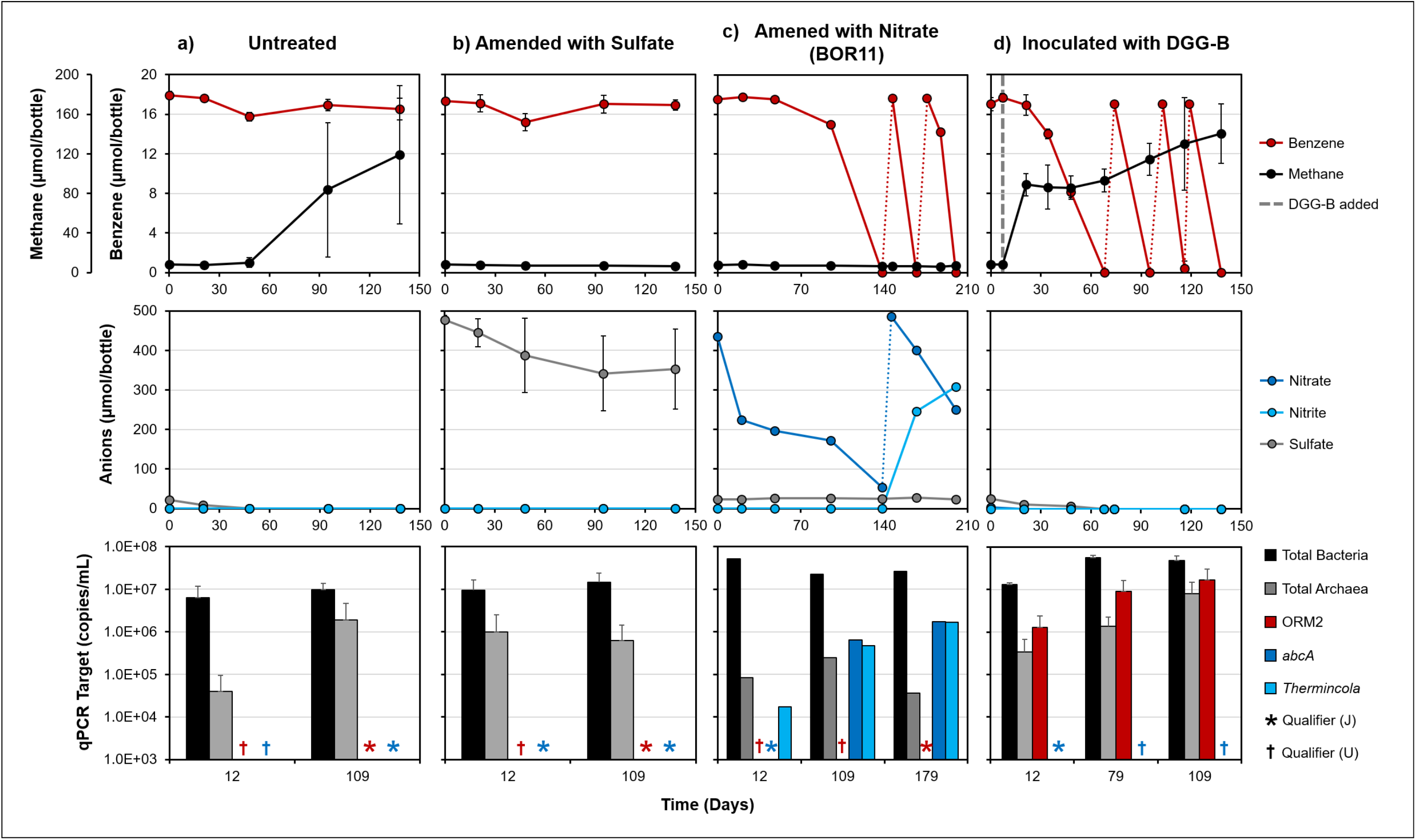
Benzene (top panels) and anion (center panels) degradation profiles of active bottles that were a) left untreated, b) amended with sulfate, c) amended with nitrate, or d) inoculated with 2.5% *v/v* DGG-B culture. Electron donor refeeding events are marked with dotted lines. The bottom panels summarize the abundance of target 16S rRNA gene copies and *abcA* enriched in each treatment. qPCR targets below quantifiable limits (< 10^3^ copies/mL) or below detection are designated by J and U qualifiers, respectively. Most results shown are the average of triplicate replicates (error bars = ± standard deviation). One replicate is shown for nitrate amended bottles; data for all replicates are shown in Figure S3. Data for bottles amended with both DGG-B and sulfate are shown in Figure S4.

Benzene was degraded in two out of three microcosms amended with 2 mM nitrate (BOR11 and BOR12 but not BOR13) within 312 days of incubation (Figures 2c and S3). Benzene depletion in the most active replicate (BOR11) was first observed after ∼ 95 days at an initial rate of 1.8 µM/day, which increased to 4.1 µM/day after re-amending benzene twice (Figure 2c). While nitrate reduction was observed throughout the incubation, near-stoichiometric amounts of nitrite were produced solely during periods of active benzene biodegradation, characteristic of incomplete nitrate reduction.^40^ Active bottles were refed nitrate once (Day 146) before clear evidence of benzene biodegradation and nitrite accumulation was observed. In benzene-degrading bottles, *abcA* gene copies increased by 3 to 4 orders of magnitude between Days 12 and 179 (Figures 2c and S3). Increases in *abcA* always coincided with the first observation of benzene loss, and *abcA* gene concentrations of least 10^5^ copies/mL appeared to be required before benzene loss was measurable. This is best seen in BOR12 (Figure S3). This bottle, which exhibited a longer lag period than BOR11, had fewer *abcA* gene copies by Day 109 (1.1 × 10^4^ copies/mL). In BOR13, where no benzene degradation activity was observed, no quantifiable amounts of *abcA* were ever detected (Figure S3). Since all known organisms that harbour an *abcA* gene belong to *Thermincola*,^15, 16, 21, 33^ gDNA samples were also screened for members of this genus. Expectedly, significant increases of *Thermincola* qPCR targets were observed in parallel with *abcA* and in similar amounts (Figures 2c and S3). The *Thermincola* qPCR primers do not exclusively capture benzene-degrading strains (see Table S3), which may explain why *Thermincola* was detected in BOR13 on Day 12 (Figure S3).

Benzene was degraded within ∼ 34 days in all bottles bioaugmented with the methanogenic consortium DGG-B (Figure 2d). The inoculum contained approximately 1.7 × 10^8^ copies/mL of deltaproteobacterial Candidate Sva0485 organisms (ORM2). We recovered 1.5 × 10^6^ copies/mL of ORM2 in bottles post-bioaugmentation which is ∼ 1% of the inoculum, within qPCR error of the 2.5% inoculum added. ORM2 gene copies increased by one order of magnitude by Day 109, supporting microbial growth post-bioaugmentation. Concentrations of archaea also increased proportionally with ORM2 (Figure 2d) and electron balance calculations confirmed that benzene oxidation was coupled to methanogenesis (Table S4). Methanogenic benzene biodegradation was also supported by the results of 16S rRNA gene amplicon sequencing (detailed below). Initial (2.0 ± 0.3 µM/day) and maximum (6.5 ± 0.3 µM/day) benzene biodegradation rates in BOR14-16 were comparable to rates reported under nitrate-reducing conditions (Table S1, Figure 2d), but benzene depletion was achieved more than 70 days sooner in bioaugmented bottles as a result of shorter lag times.

The addition of 2 mM sulfate to bioaugmented bottles (BOR17-19) resulted in no significant differences in benzene lag times, degradation rates, or enrichment of deltaproteobacterium ORM2 relative to bioaugmentation only (Table S1, Figure S4). Electron balances link benzene oxidation primarily to methanogenesis, although some benzene oxidation appears to have been linked to sulfate reduction (Table S4). Electron balance calculations revealed an unknown source of electron acceptor demand from the BOR soil that was more abundant than the hydrocarbons added, which confounded electron balances in some cases. Given that all benzene-degrading members of Sva0485 have been characterized as fermentative organisms (Figure 1),^17, 20, 30^ sulfate may not enhance benzene degradation by members of this candidate clade. Microbial community sequencing (described later) offered additional clues as to the microorganisms and metabolisms driving benzene biodegradation in these bottles.

### 3.2 Effect of Hydrocarbon Co-Contaminants on Anaerobic Benzene Bioremediation

We next explored the impact of TEX and naphthalene on benzene biodegradation (bottles BOR20-38). We observed variable patterns of BTEX degradation and lag times among replicates and treatments, although naphthalene was recalcitrant in all cases (Figures S5–S10). Toluene was the only hydrocarbon degraded in untreated bottles (BOR24-26), although degradation stalled in 2 out of 3 replicates after sulfate was depleted (Figure S6). The addition of 2 mM sulfate (BOR27-29) enhanced the biodegradation of toluene, *o*- and *m-*xylene (Figure S7), whereas 2 mM nitrate (BOR30-32) stimulated complete BTEX degradation in two out of three microcosms (Figure S8), and degradation proceeded in the following order: toluene > *m*-xylene > ethylbenzene > *o*-xylene > benzene. Perhaps as a result of being degraded last, benzene lag times increased from 95-160 days (in BOR11-12) to 170-260 days in BOR31-32 (Table S1). Initial rates in BOR31-32 were 17-72% slower than rates reported for BOR11.

In DGG-B bioaugmented bottles (BOR33-35), toluene, benzene and *o*-xylene were degraded in roughly the same period of ∼ 160 days (Figure S9). Looking more closely, it is apparent that toluene degradation was initiated quickly depleting the available sulfate, stopped when sulfate was depleted, and resumed when methanogenic conditions became established. In the bioaugmented bottles BOR36-38 that received additional sulfate, toluene was degraded much more rapidly (<40 days), as were *o*- and *m*-xylene, while benzene degradation began after a similar lag as BOR33-35 (Table S1, Figure S10). Initial rates of benzene degradation were slower than in benzene-only bottles. We re-amended BTEX and naphthalene to sulfate-amended bottles BOR37 and BOR38 on Day 146 and found that benzene degradation rates increased from ∼ 0.6 µM/day to ∼ 1.4 µM/day (Figure S10), still slower than initial rates reported for BOR14-19. We refed BOR38 a third time with benzene only and this time degradation rates increased up to 4.8 µM/day (Figure S10). In bottles BOR36-38, electron balance calculations revealed that benzene degradation became coupled to sulfate reduction, as much more benzene was consumed than could be explained by observed methane production (Table S5). Ethylbenzene and naphthalene were not degraded in these treatments.

### 3.3 Rates of Anaerobic Benzene Biodegradation and Microbial Growth Estimates

We next examined the relationship between rates of benzene degradation and the abundance *abcA/Thermincola* and Sva0485 clade Deltaproteobacteria as measured by qPCR. An estimate of the zero order rate of benzene degradation (µmoles per L/day) was calculated for each time interval where corresponding qPCR data was also available. A good correlation between benzene degradation rate and microbial abundance was observed above a threshold between 10^4^ or 10^5^ cells/mL for both targets (Figure 3). Below this threshold, benzene degradation was very slow or not detected. In the presence of TEX and naphthalene, *abcA* and ORM2 copies/mL increased less than in bottles containing benzene only, and corresponding rates of anaerobic benzene biodegradation were proportionally slower (Figure 3). Extrapolating a linear regression from both datasets, we calculated estimates of the concentration of *abcA* (3.8 × 10^5^ copies/mL) and ORM2 (4.3 × 10^6^ copies/mL) required to achieve a rate of 0.1 µM benzene/day (∼ 8 µg/L/day). From these data, about an order of magnitude fewer *abcA*-containing organisms are required compared to ORM2. Overall, these data suggested that benzene biodegradation was partially impeded by the hydrocarbon co-contaminants themselves or as a result of their degradation, leading to slower benzene removal and lower abundance of benzene-degrading microbes.

**Figure 3.**
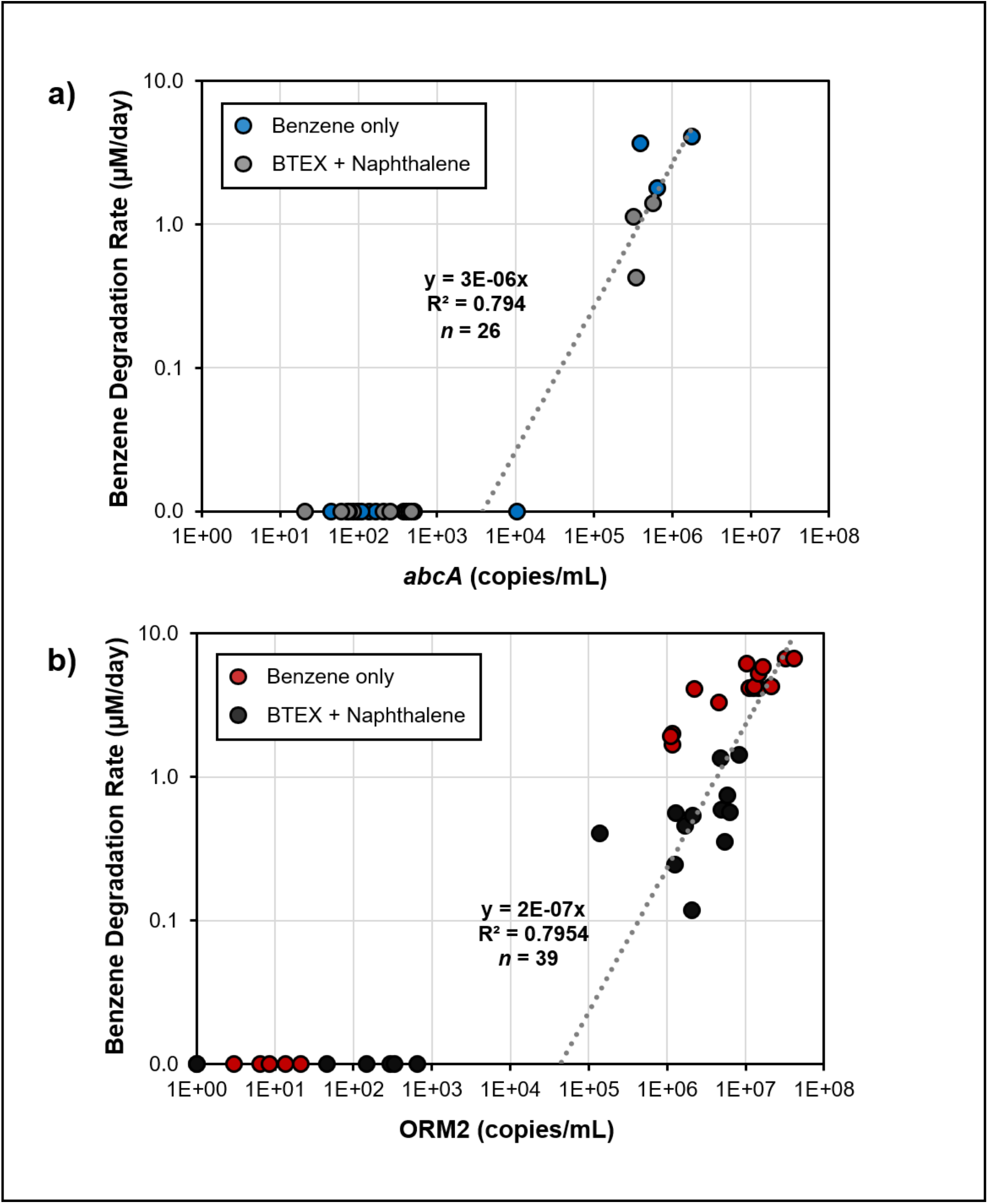
qPCR data showing correlations between concentrations of a) *abcA* or b) Sva0485 clade Deltaproteobacteria and rates of anaerobic benzene degradation. Samples are grouped by substrate. Day 12 gDNA samples were omitted from correlation analyses as rate data were inacurate so early in the study.

We used qPCR and concentration data to estimate doubling times and specific yields (Table S5). Although the data are relatively sparse, the fastest doubling times were observed under nitrate-reducing conditions (10 ± 2 days; *n*=6). Yields were estimated at 7 ± 2 × 10^3^copies *abcA*/nmol benzene. In bottles bioaugmented with DGG-B, doubling times were 20 ± 2 days (*n*=8) in bottles containing benzene only. In bottles with BTEX and naphthalene, doubling times for ORM2 were longer and more variable (50 ± 10 days; *n*=6). Yield estimates for ORM2 were similar regardless of treatment at about 3 ± 1 × 10^5^ copies ORM2/nmol benzene. ORM2 cell are smaller than *Thermicola* cells, and when mass per cell is taken into account, yields are not so dissimilar (∼0.05 to 0.07 g cells per g benzene; Table S6). These doubling times are remarkably consistent with reported doubling times of 9 days (nitrate-reducing) and 30 days (methanogenic) reported as far back as 2003.^25^ Luo *et al*.^20^ reported doubling times for the OR consortium in 2016 of 34 days and a yield for ORM2 of about 0.01-0.02 cells/g benzene. These yields are very low.

### 3.4 Microbial Community Analysis

Eighty-six (86) gDNA samples from active microcosms were analyzed by 16S rRNA gene amplicon sequencing, including the original BOR sediments and the DGG-B inoculum. Table S7 provides results for amplicon sequence variants (ASVs) with greater than 0.1% relative abundance in at least one sample. Non metric multidimensional scaling (NMDS) ordination identified three clusters based on bioremediation treatment (Figure S11). Communities enriched in untreated and sulfate-amended bottles were comparable to each other, and were most closely related to original BOR sediments. All bioaugmentation bottles clustered together, and were more similar to the DGG-B inoculum. Nitrate biostimulation enriched for distinct communities. Microbial communities from bottles amended with mixed hydrocarbons did not cluster distinctly from their counterparts amended with benzene only (Figure S11).

A heatmap comparing the microbial communities in all bottles after 109-258 days of incubation (Figure 4) recapitulates the three main clusters seen in Figure S11. The first cluster regroups untreated and sulfate amended bottles, as well as the original Borden sediments. The latter were predominantly comprised of Betaproteobacteria (68.8%, collectively), including *Hydrogenophaga* and unclassified *Burkholderiaceae*. Sulfate-amended treatments became enriched in Deltaproteobacteria, in particular *Geobacter* (3.35 – 14.4%) and *Desulfovibrio* (0.83 – 24.7%). Benzene degradation was not observed in any of these bottles. We did detect enrichment of a sequence variant (*Geobacter* ASV3) with 97.4% sequence identity to *Geobacter metallireducens* strain Ben, an iron-reducing isolate previously shown to degrade benzene,^26, 41^ but no other predicted anaerobic hydrocarbon degraders were identified. Dominant ASVs enriched in the presence of BTEX and naphthalene were comparable to those enriched on benzene alone (Figure 4).

**Figure 4.**
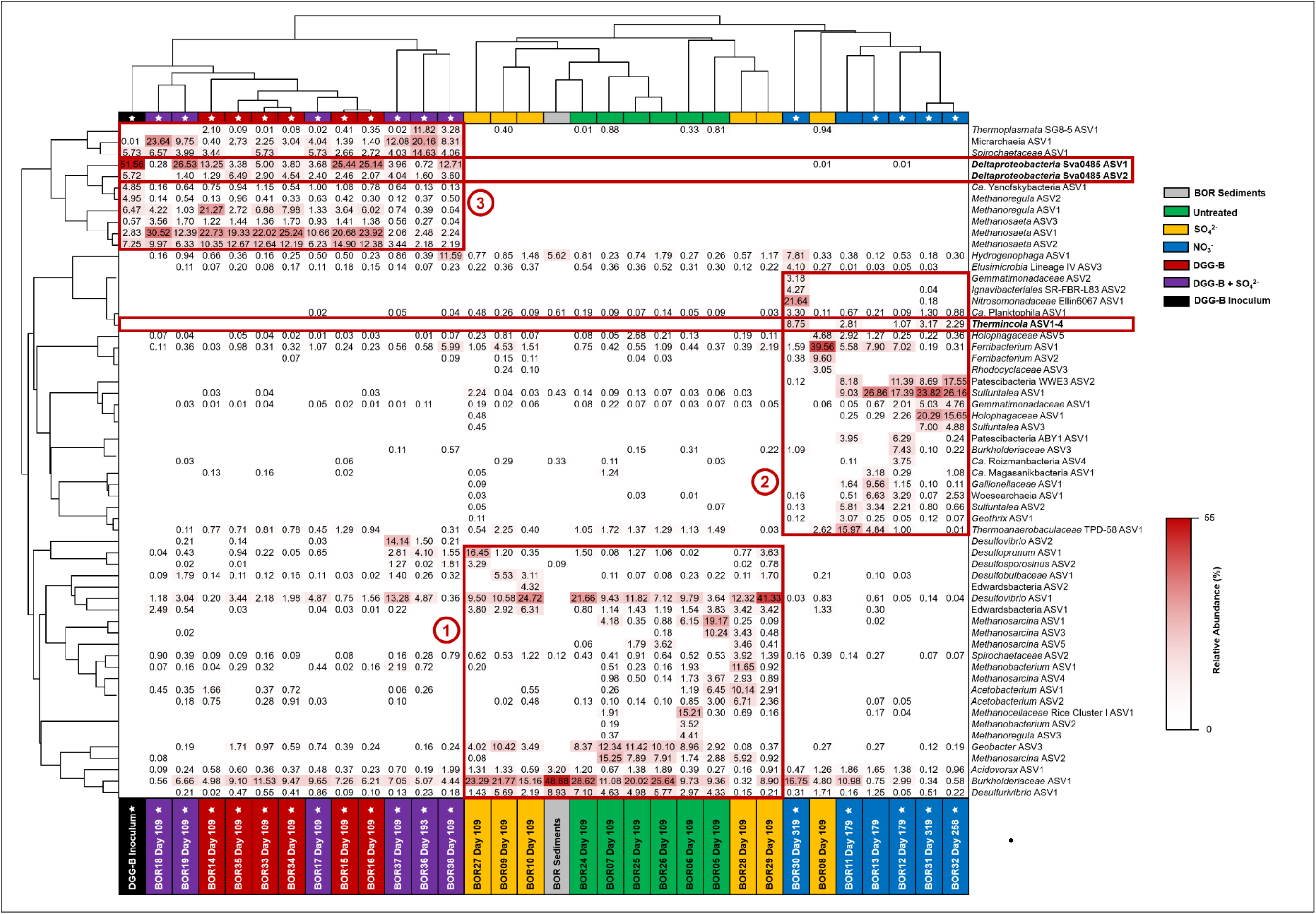
**(above):** Microbial community analysis of dominant ASVs (≥ 3% in one or more sample) in experimental bottles at select timepoints, relative to homogenized CFB Borden sediments and the DGG-B inoculum. Samples marked with stars (★) indicate active anaerobic benzene degradation at the time of sampling. Rows and columns were clustered in ClustVis^55^ using correlation distance and average linkages. The inset heat map designates the relative abundance of each ASV, where darker shades of red indicate higher relative abundances. The relative abundance of minor ASVs is provided in Table S7. Predicted benzene-degrading bacteria and microbial clusters of interest are highlighted in red boxes. Finally, four *Thermincola* ASVs have been collapsed and included in this figure to highlight their enrichment in active nitrate-reducing, benzene-degrading bottles.

The second cluster regroups all nitrate-amended bottles and one bottle (BOR08) amended with sulfate that exhibited much less sulfate reduction compared to other bottles of the set (Table S4). Twelve unique ASVs belonging to *Thermincola* ASVs were identified across our dataset, but four (designated as *Thermincola* ASV1-4) were enriched (up to 8.8% of reads) solely in active nitrate-reducing, benzene-degrading replicates (BOR11-12 and BOR 30-32; Table S7, Figures 4 and S12a). The relative abundance of these four *Thermincola* ASVs was nearly identical across all samples, suggesting they all belong to the same organism and genome. *Thermincola* ASV1-4 shared 97.4 – 99.2% identity to sequences obtained from benzene degrading nitrate-reducing enrichment cultures in our lab (Figure S13).^21, 34, 35^ The four ASVs also shared high sequence identity (94.7 – 98.3%) to several other *Thermincola* species highlighted in Figure 1. Given that the enrichment of *Thermincola* 16S rRNA gene copies coincided with that of *abcA*, and that expression of *abcA* has previously been demonstrated at CFB Borden,^32^ we propose that *Thermincola* is likely a key anaerobic benzene degrader at this field site. Another ASV that was enriched solely in active nitrate-reducing microcosms belonged to Patescibacteria candidate division WWE3 (Table S7, Figures 4 and S12a). Members of the Patescibacteria superphylum (referred to as the Candidate Phyla Radiation, or CPR) have been identified in benzene-degrading communities, and are also prevalent in groundwater, sediment, lake, and other aquifer environments.^42^ They have tiny genomes and exhibit fermentative communal lifestyles.^43^ Our data hints that benzene-contaminated groundwater may enrich for select members of this expansive phylogenetic group of candidate organisms.

The third cluster includes all DGG-B bioaugmented microcosms (BOR14-19, BOR33-38), where a limited core of ∼ 11 ASV were enriched by Day 109 (up to 79.1% of reads), all of which likely originated from the inoculum (Table S7, Figures 4 and S12b). Two sequence variants affiliated with Sva0485 clade *Deltaproteobacteria* (ORM2) and an ASV associated with *Candidatus* Yanofskybacteria (previously referred to as Parcubacterium/OD1) were the most abundant bacteria. High abundances of *Ca*. Yanofskybacteria have previously been reported in the OR consortium (now DGG-B), although its specific community role is not understood.^20^ Methanogenic archaea belonging to *Methanoregula* (2 ASVs) and *Methanosaeta* (3 ASVs) all increased in relative and absolute abundances over time (Figure S12b), agreeing with qPCR trends reported in this study and in previous studies.^20^ *Deltaproteobacteria* Sva0485 ASV2 is 100% identical to both deltaproteobacterium ORM2a and ORM2b reported previously,^20^ whereas Sva0485 ASV1 differed by a single nucleotide position (Figure S14). The significance of this second sequence variant (different strain, mutation or error) is not clear at this time. Its high sequence abundance and presence in all bioaugmented samples supports that Sva0485 ASV1 is not a sequencing artifact. We also observed growth of *Spirochaetaceae* (Table S6, Figures 4 and S12b), a family previously hypothesized to participate in recycling of dead biomass in anaerobic hydrocarbon-degrading communities.^44^ Other major organisms in the third cluster include Micrarchaeia and an unclassified Thermoplasmata not previously detected in the DGG-B inoculum. These are archaeal clades whose poorly characterized members also have tiny genomes that have eluded recognition in the past. In this study, Micrarchaeia were more enriched in sulfate-amended bottles (Figure 4). *Desulfovibrio* ASV2 was also enriched in sulfate-amended bottles, particularly those amended with the suite of BTEX hydrocarbons (BOR36-38). This ASV did not cluster with others from the DGG-B inoculum, however, so it may have originated from the sediment.

Reconciling the microbial community profile with qPCR data, benzene biodegradation rates and electron balance data, we see tremendous concordance. While electron acceptors drove differences in the microbial communities more so than electron donors, we clearly see proliferation of specific benzene degrading organisms only where benzene degradation was observed. Organisms that proliferate in concert with predicted benzene degraders provide clues to essential syntrophic partners. In *Thermincola* cluster 2 (Figure 4), we see enrichment of *Bulkholderiaceae* and *Sulfuritalea*; both implicated in degradation of aromatics such as benzoate with nitrate as electron acceptor.^45^ *Ferribacterium* from the family of Rhodocyclaceae also became enriched in Cluster 2 particularly in the oddball bottle BOR08. Although not measured, it might have reduced iron in the BOR sediments coupled to oxidation of hydrogen or organic acids, explaining lower sulfate reduction for this bottle. Interestingly, other benzene-degrading *Thermincola* have been characterized from iron-reducing cultures. In cluster 3 (Figure 4), proliferation of a narrow group of microbes surrounding Sva0485 clade Deltaproteobacteria (ORM2) was evident, and perhaps the tight association, particularly with *Candidatus* Yanofskybacteria and Methanosaeta was disrupted by other hydrocarbon degraders when TEX and naphthalene were added to the bottles in addition to benzene.

## 5 Perspectives for the Field

Nitrate addition was found to stimulate benzene degradation and growth of *abcA*-containing *Thermincola* as well as degradation of TEX co-contaminants (Figure S8), although replicates were not consistent. Several *in situ* studies have also reported successful BTEX attenuation using nitrate amendment,^3,4, 46^ thus nitrate biostimulation does appear to offer site managers an option for field remediation. Nitrate and nitrite are themselves regulated groundwater contaminants and repeated nitrate amendment will be necessary to sustain activity, as was the case in our tests (Figures 2c, S3 and S8). Thus, nitrate addition may not be a practical or cost-effective solution for all sites.

Bioaugmentation with methanogenic consortium DGG-B resulted in growth of the entire benzene-degrading community and sustained biodegradation (Figures 2d, 4, S4, S9, S10 and S12b). Bioaugmentation is not thought to be needed for hydrocarbons because these substrates occur naturally and indigenous degraders are thought to be ubiquitous in nature.^47, 48^ Moreover, many sites already have ample sources of anaerobic electron acceptors (nitrate, sulfate, and/or CO_2_) that should support benzene-degrading communities. This begs the question why do intrinsic benzene-degrading organisms fail to become enriched in anaerobic environments? Especially since microbes that metabolize toluene and xylenes anaerobically proliferate far more commonly?

One hypothesis is that growth of anaerobic benzene degrading organisms is inhibited by the presence of other hydrocarbons.^3, 49, 50^ Such a hypothesis is somewhat consistent with the data from this study (Figures 3, S8-S10). Average rates of anaerobic benzene degradation decreased in bottles containing co-contaminants. Although inhibition due to toxicity is one explanation,^51, 52^ it is unlikely since background methanogenesis and sulfate reduction were not impacted. Competition for essential nutrients or co-factors, predation by other organisms, cross-feeding, leaky pathways, and high decay rates are among many other more likely explanations. Given the co-enrichment of a narrow group of 11 ASVs in cluster 3 (Figure 4), it seems that there are very significant co-dependencies between the key benzene-degraders, members of the CPR and specific methanogens. Further experimentation is required to narrow down these possibilities.

It is now clear that microorganisms responsible for anaerobic benzene transformation are highly specialized, occupying a tiny single-substrate niche, akin to how certain *Dehalococcoides* are uniquely adapted to only a few chlorinated substrates. ^53^ We have been unable to identify any other substrate other than benzene for *Thermincola* nor for Sva0485 clade Deltaproteobacteria, and none of the genes in their genomes provide further hints. Amending an intermediate compound like benzoate simply enriches for other organisms in the community and thus does not promote their growth.^21, 31^ Bioaugmentation with specialized anaerobic dechlorinating bacteria at sites contaminated with chlorinated solvents has shown tremendous success for over 20 years as reviewed in Stroo *et al*.^54^ Perhaps we could tackle benzene in anoxic environments in a similar way. To this end, we are evaluating DGG-B bioaugmentation in an increasing number of laboratory microcosms and to date have initiated three bioaugmentation field pilots. Though it remains unclear why intrinsic anaerobic benzene degraders rarely proliferate *in situ*, we are encouraged to determine if bioaugmentation to boost microbial community abundance could be effective for BTEX cleanup in anoxic environments.

## 6 AUTHOR CONTRIBUTION

EE, FL and SD contributed to the conception of this research project. DGG-B culture maintenance was overseen by all authors. FL, JW and NB established, maintained and sampled all microcosms during treatability testing. Molecular sampling and testing was conducted by CT, FL, SG and NB. CT processed and interpreted all molecular results and drew figures. CT and EE wrote the manuscript.

## 7 ASSOCIATED CONTENT

The Supporting Information is available free of charge of the ACS Publications website. Supporting Information includes detailed description of analytical methods, and DNA extraction and associated PCR and amplicon analyses (PDF). The Supporting information PDF file also includes 14 supplemental figures showing various plots of concentration data over time for each treatment, as well as qPCR calibration data and sequence alignments. A supplementary excel file includes all Tables (S1-S7) providing experimental design, raw and transformed data, electron balances, yield and doubling time calculations, and a table of amplicon sequence data.

## 8 ACKNOWLEDGEMENTS

This study was funded by a Genomic Application Partnership Program (GAPP) grant awarded to Elizabeth Edwards and Sandra Dworatzek (Project ID OGI-102), which was supported by Genome Canada, Genome Ontario, the Government of Ontario, Mitacs Canada, SiREM and Federated Co-Operatives Limited. Additional financial support was provided through a Natural Sciences and Engineering Research Council (NSERC) Discovery grant awarded to EE. The authors gratefully thank the laboratory of Dr. Neil Thomson (University of Waterloo) for field sample collection, and Dr. Camilla Nesbø (University of Alberta) for assistance with Illumina sequencing analysis.

---

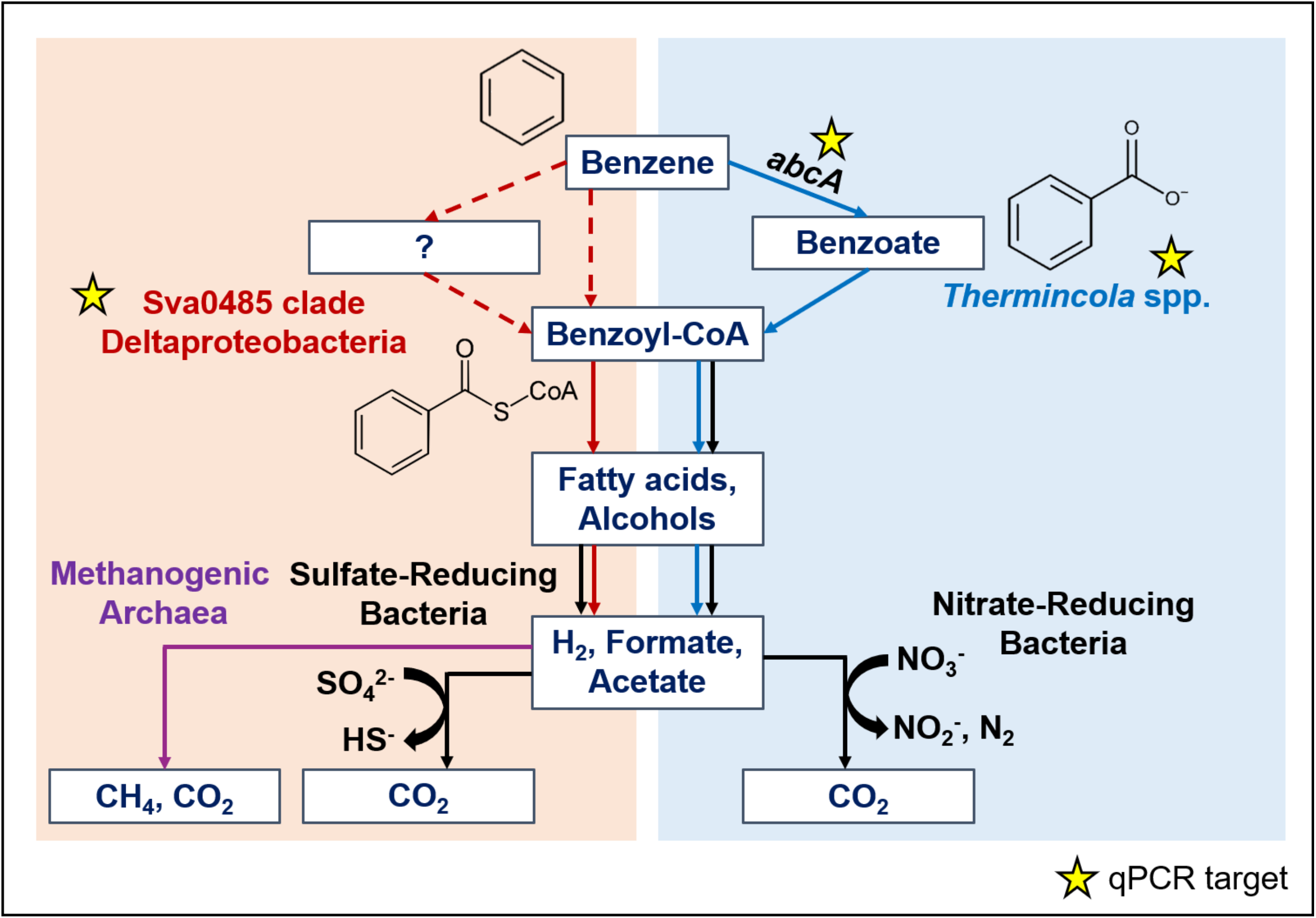

