## Supplemental Material for "Anaerobic Benzene Biodegradation Linked to Growth of Highly Specific Bacterial Clades"

Number of Pages: 21

Number of Supporting Texts: 4

Number of Tables: 7 (all except Table S1 in accompanying Excel file)

Number of Figures: 14

### TABLE OF CONTENTS


#### Supporting Texts

**Text S1:** Description of Figure 1 maximum likelihood phylogenetic tree construction

**Text S2:** Detailed analytical methods for monitoring hydrocarbon and anion concentrations

**Text S3:** Standard operating procedures and calibration information for all qPCR assays

**Text S4:** 16S rRNA Amplicon Sequencing method and analysis details

#### Supplementary Tables (all except Table S1 are in accompanying Excel file)

**Table S1.** Experimental treatments and overview of results (this document and Excel file)

**Table S2 (a&b).** Summary of all microcosm amendment and sampling events, including concentrations for hydrocarbons, anions and qPCR targets (Table S2a: concentration in mg/L; Table S2b: concentration in µmol/bottle)

**Table S3:** Primers used for quantification of targeted 16S rRNA and *abcA* genes

**Table S4:** Electron balances in active microcosms amended with benzene

**Table S5:** Electron balances in active microcosms amended with BTEX and naphthalene

**Table S6:** List of all gDNA samples amplified by Illumina sequencing in this study, including the percent abundance of ASVs $\geq$ 0.1% abundance and their taxonomic assignment

**Table S7:** Growth yield calculations (copies per nmol benzene consumed) and estimates of doubling times for *abcA* and deltaproteobacterium ORM2

#### Supplementary Figures

**Figure S1:** Standard curves and efficiencies of qPCR runs to quantify specific microbial groups

**Figure S2:** Benzene and anion degradation profiles of benzene-amended mercuric chloride-poisoned sterile control bottles and active microcosms that were left untreated or amended with 2 mM sulfate

**Figure S3:** Benzene and anion degradation profiles of active bottles amended with 2 mM nitrate

**Figure S4:** Benzene and anion degradation profiles of active bottles amended with 2.5% *v/v* DGG-B culture and 2 mM sulfate

**Figure S5:** Mixed hydrocarbon and anion degradation profiles of mercuric chloride poisoned sterile control microcosms

**Figure S6:** Mixed hydrocarbon and anion degradation profiles of active microcosms that were left untreated

**Figure S7:** Mixed hydrocarbon and anion degradation profiles of active microcosms amended with 2mM sulfate

**Figure S8:** Mixed hydrocarbon and anion degradation profiles of active microcosms amended with 2 mM nitrate

**Figure S9:** Mixed hydrocarbon and anion degradation profiles of active microcosms inoculated with 2.5% *v/v* DGG-B culture

**Figure S10:** Mixed hydrocarbon and anion degradation profiles of active microcosms inoculated with 2.5% *v/v* DGG-B culture and amended with 2 mM sulfate

**Figure S11:** Bray-Curtis NMDS plot of all 16S rRNA gene amplicon samples

**Figure S12**. Time course microbial community composition of a) nitrate biostimulation microcosm BOR11 and b) DGG-B bioaugmentation microcosm BOR16, relative to original Borden sediments and DGG-B inoculum.

**Figure S13:** Multiple sequence alignment of *Thermincola* ASV1-4 against two reference 16S rRNA gene sequence clones from nitrate-reducing, benzene-degrading *Thermincola* spp.

**Figure S14:** Multiple sequence alignment of *Deltaproteobacteria* Sva0485 ASV1-2 against reference 16S rRNA gene sequence clones for *Deltaproteobacteria* ORM2a and ORM2b.

**References for SI**

### Supporting Information Text S1:

#### Description of Figure 1 maximum likelihood phylogenetic tree construction, and 16S rRNA gene sequences of predicted anaerobic benzene degraders without available accession numbers

The final consensus maximum likelihood phylogenetic tree presented in Figure 1 was constructed in MEGA7 using the Tamura-Nei distance method and performing 500 bootstrap replicates. Initial tree(s) for the heuristic search were obtained automatically by applying Neighbor-Join and BioNJ algorithms to a matrix of pairwise distances estimated using the Maximum Composite Likelihood (MCL) approach, and then selecting the topology with superior log likelihood value. The tree was drawn to scale, with branch lengths measured in the number of substitutions per site. Codon positions included were 1st+2nd+3rd+Noncoding. All nucleotide positions with less than 95% site coverage were eliminated. That is, fewer than 5% alignment gaps, missing data, and ambiguous bases were allowed at any position. The anaerobic benzene-degrading archaeon *Ferroglobus placidus* was used to root the tree.[1] Bootstrap values < 60% are not shown. We verified the taxonomy of each uncultured benzene-degrading strain using the Silva 138 small subunit (SSU) rRNA database.[2] The 16S rRNA gene sequence for the enrichment culture BPL *Pelotomaculum* clone is published in Dong et al.[3]

**Text S2: Analytical methods for monitoring hydrocarbon and anion concentrations**

Methane, BTEX, and naphthalene dissolved in the aqueous phase of the microcosms were measured using an Agilent 7890 gas chromatograph (GC) equipped with an Agilent G1888 headspace autosampler (see Supplementary Materials). Liquid samples (500 µL) were injected into autosampler vials containing 5.5 mL of acidified deionized water (pH ~ 2). See method details in SI. The water was acidified to inhibit microbial activity between microcosm sampling and GC analysis. Vials were sealed with an inert Teflon-lined septum and aluminium crimp cap for automated injection of 3 mL of headspace onto the GC. The autosampler was programmed to heat each sample vial to 75°C for 45 min prior to headspace injection (3 mL) into a GSQ Plot column (0.53 mm × 30 m) and a flame ionization detector. Vial heating ensured that all volatile organic compounds in the aqueous sample would partition into the headspace. The injector temperature was 200°C, and the detector temperature was 250°C. The oven temperature was programmed as follows: 35°C for 2 min, increased to 100°C at 50 °C/min), then increased to 185°C at 25°C/min and held at 185°C for 6.80 min. The carrier gas was helium at a flow rate of 11 mL/min. Calibration was performed using external standards purchased as standard solutions (Sigma, St Louis, MO), where known volumes of standard solutions were added to acidified water in auto sampler vials and analysed as described above for microcosm samples.

Anion (nitrate, nitrite, sulfate, acetate equivalents, chloride, and phosphate) analysis was performed on a Thermo-Fisher ICS-2100 ion chromatograph (IC) equipped with a Thermo-Fisher AS-DV autosampler, an AS18 column and a sample loop volume of 25 µL. An isocratic separation was performed using 33 millimolar (mM) reagent grade sodium hydroxide eluent generator cartridge (Thermo Scientific, Burlington, ON) eluent for 13 min. External standards were prepared gravimetrically using chemicals of the highest purity available (Sigma, St Louis, MO or Bioshop, Burlington, ON), and analysed with each set of samples. IC samples were prepared by centrifuging (13,000 rpm for 5 min) 250 µL of culture, diluting the supernatant 50-fold in deionized water, and placing the final volume in a Thermo-Fisher autosampler vial with a cap that filters the sample during automated injection onto the IC. pH measurements (Oakton pH Spear, Vernon Hills, IL) and anion data for acetate equivalents, chloride and phosphate are not discussed in the main text, but are provided in Table S2 (raw data).

Electron balances provided in Tables S4 and S5 were calculated from the data in Table S2b to constrain relationships between hydrocarbon electron donors and available electron acceptors. No significant changes in hydrocarbon or anion concentrations were observed in mercuric chloride-poisoned sterile control bottles. As stated in main text, background acceptor demand from the Borden soil itself was substantial, as seen in Tables S4 and S5 (Acceptor-Donor column). In nitrate-amended bottles, nitrate was clearly the acceptor for hydrocarbon degradation. In bioaugmented bottles without added sulfate, methanogenesis could account for all hydrocarbon depleted. In bioaugmented bottles with sulfate added, both sulfate reduction and methanogenesis were observed (Tables S4 and S5). Three bottles (BOR04, BOR18, and BOR28) were found to have leaky caps, which caused some abiotic hydrocarbon losses during the incubation period. The leaky caps also allowed for hydrogen ingress (from the glovebox atmosphere), allowing for electron acceptor consumption and/or methane production (Tables S4 and S5). No substantial changes to hydrocarbon or anion concentrations were reported after caps were replaced. The methane data reported in this study is highly inaccurate (affecting electron balances only) largely because analysis was based on dissolved concentrations measured in aqueous samples collected for all analyses, while most of the methane would be in the headspace.

**Text S3: Standard operating procedures and calibration information for all qPCR assays**

Amplification of targeted genes was carried out in 20 µL reactions containing 500 nM of each designated forward and reverse primer (see Table S3), 2 µL template DNA, 10uL 2 × SsoFast™ EvaGreen® Supermix (Bio-Rad Laboratories, Hercules, CA), and UltraPure™ DNase/RNase-Free Distilled Water (Invitrogen, Carlsbad, CA). Serial dilutions of plasmids containing corresponding targeted gene fragments were used to generate standard curves. Each qPCR reaction was performed in duplicate on a Bio-Rad CFX96 real time PCR machine. Thermal cycling conditions were as followed; an initial denaturation step (2 min at 98°C), 40 cycles of 5 s at 98°C and X °C (X = 59°C if targeting *abcA*, ORM2, GenArch or PeptoBen primers; 55°C if using GenBac primers), and melt curve analysis (65–95°C with an increase of 0.5°C every 10 s). qPCR results were analyzed using Bio-Rad CFX Manager software. The amplification efficiency for each qPCR run batch range from 87% to 105% (GenBac, 96.6%; GenArch, 90.2%; ORM2, 95.4%; *abcA*, 97.9%; PeptoBen, 92.8%), with $R^2$ values of >0.995.

**Text S4: 16S rRNA Amplicon Sequencing method and analysis details**

The relative abundance of other microbes was determined using 16S rRNA gene amplicon sequencing. Amplification and Illumina sequencing (MiSeq 300PE; paired-end) of extracted gDNA was carried out by McGill University at the Genome Quebec Innovation Centre. All amplicon sequence reads were generated using modified, "staggered end" primers 926F (AAACTYAAAKGAATWGRCGG) and 1392R (ACGGGCGGTGWGTRC), where 0 – 3 random bases were inserted between the primer and Illumina adaptor sequences. Creating staggered ends improves sequencing quality.[4] Read processing and sequencing analyses were performed in QIIME 2 version 2019.10.[5] Raw reads were trimmed of primer sequences and staggered ends, truncated, and denoised using the DADA2 pipeline within QIIME 2.(Callahan et al., 2016) Quality filtered forward (260 bp) and reverse (240 bp) reads were then merged with a maximum of 2 expected errors in the overlap region. Chimeric sequences and sequences < 400 nucleotides in length were removed. The final amplicon sequence variants (ASVs) were classified against the SSU SILVA 132 database[2] trained on the targeted 16S rRNA gene region. Non-metric multidimensional scaling (NMDS) ordination using Bray–Curtis distance metrics was used to visualize overall grouping patterns of ASVs. We also used ClustVis[6] to prepare correlation distance and average linkage heatmaps of dominant ASVs in select gDNA samples. Raw sequence reads were deposited to the National Center for Biotechnology Information Short Read Archive (SRA) under BioProject PRJNA661350. No 16S rRNA gene amplicons were recovered from sterile poisoned control bottles at any timepoint.

**Table S1. Experimental treatments and overview of results**

| Experimental Treatment | Assigned Bottle Names | Replicates | Amendments | Days of Incubation | Benzene Biodegradation Reported? | Approximate Lag Period (Days)[a] | Initial Benzene Degradation Rate (µM/day)[a,b] |
|---|---|---|---|---|---|---|---|
| **Treatments containing benzene only** amended at 7 mg/L (approximately 17 µmol/bottle) | | | | | | | |
| Sterile Control | BOR01-04 | 4 | 0.05% (w/v) $HgCl_2$ | 645 | No (0/4) | N/A | N/A |
| Untreated (Natural Attenuation) | BOR05-07 | 3 | None | 645 | No (0/3) | N/A | N/A |
| Sulfate Amended | BOR08-10 | 3 | 2mM $Na_2SO_4$ | 645 | No (0/3) | N/A | N/A |
| Nitrate Amended | BOR11-13 | 3 | 2 mM $NaNO_3$ | 319 | Yes (2/3) | 95 – 140 | 1.8 – 3.0 |
| DGG-B Inoculation (Bioaugmentation) | BOR14-16 | 3 | 2.5% *v/v* DGG-B | 138 | Yes (3/3) | ~30 | 2.0 ± 0.3 |
| DGG-B + Sulfate | BOR17-19 | 3 | 2.5% *v/v* DGG-B + 2 mM $Na_2SO_4$ | 138 | Yes (3/3) | ~30 | 1.9 ± 0.2 |
| **Treatments containing BTEX + Naphthalene** amended as followed: 5 mg/L benzene, 5 mg/L toluene, 1.5 mg/L ethylbenzene, 2 mg/L *o*-xylene, 1.5 mg/L *m*-xylene, and 2 mg/L naphthalene | | | | | | | |
| Sterile Control | BOR20-23 | 4 | 0.05% *w/v* $HgCl_2$ | 312 | No (0/4) | N/A | N/A |
| Untreated (Natural Attenuation) | BOR24-26 | 3 | None | 312 | No (0/3) | N/A | N/A |
| Sulfate Amended | BOR27-29 | 3 | 2mM $Na_2SO_4$ | 312 | No (0/3) | N/A | N/A |
| Nitrate Amended | BOR30-32 | 3 | 2 mM $NaNO_3$ | 312 | Yes (3/3) | 170 – 260 | 1.0 ± 0.5 |
| DGG-B Inoculation (Bioaugmentation) | BOR33-35 | 3 | 2.5% *v/v* DGG-B | 312 | Yes (3/3) | ~40 | 0.5 ± 0.1 |
| DGG-B + Sulfate | BOR36-38 | 3 | 2.5% *v/v* DGG-B + 2 mM $Na_2SO_4$ | 312 | Yes (3/3) | 40 – 170 | 0.6 ± 0.2 |

[a]N/A = not applicable. No benzene degradation was observed.

[b]Benzene degradation rates were calculated using two timepoints per replicate; the first with clear onset of benzene consumption, and the second where benzene could no longer be detected. Error bars indicate standard deviation of triplicate bottles. Lower and upper rates are provided for treatments where benzene degradation activity was only observed in 2 replicates.

### NB: Table S1 (repeated) along with Remaining Supplemental Tables (S2-S7) are all found in the accompanying Excel File

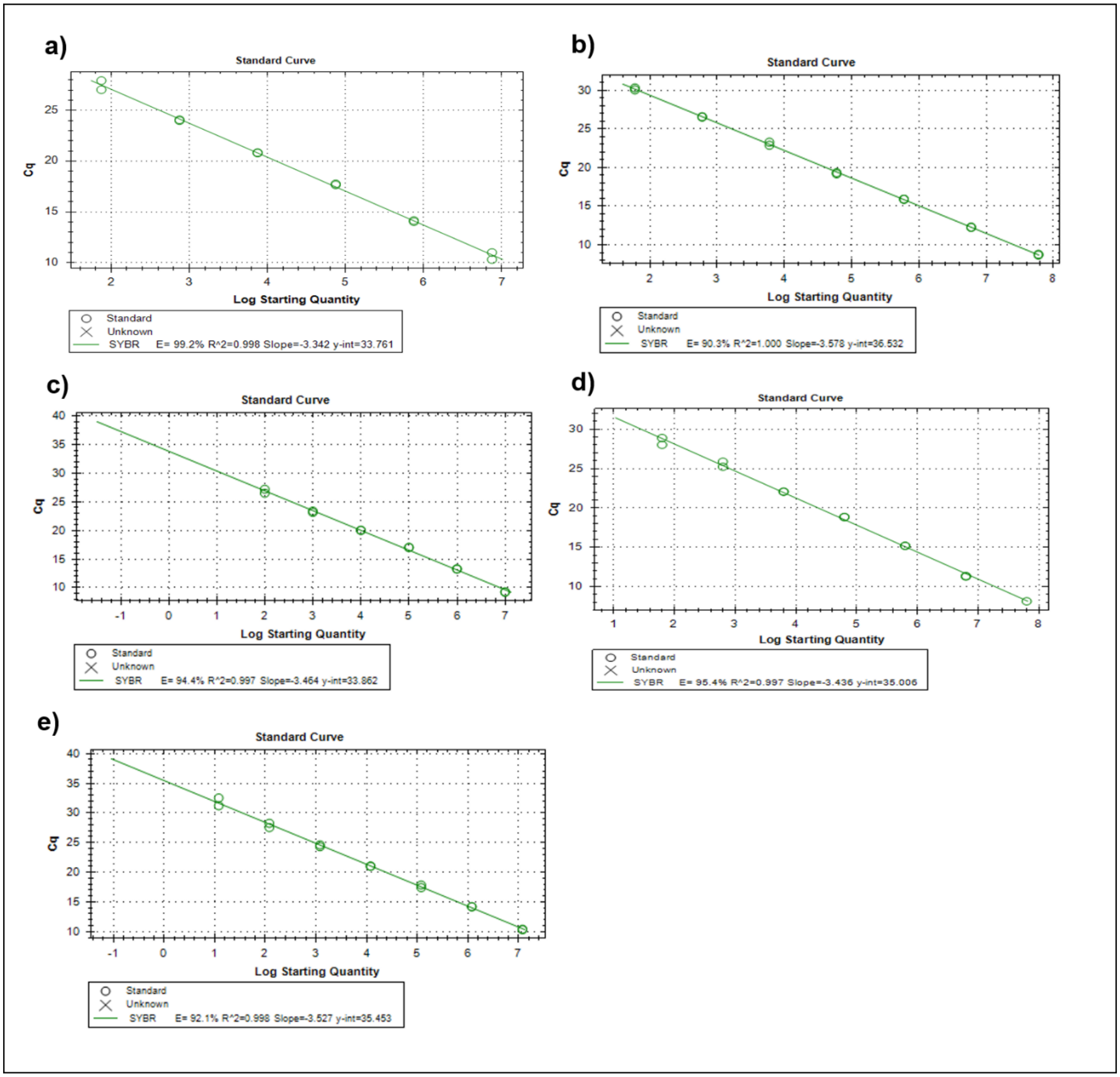


**Figure S1:** Standard curves and efficiencies of qPCR runs to quantify a) Total Bacteria, b) Total Archaea, c) deltaproteobacterial candidate clade Sva0485 (including deltaproteobacterium ORM2), d) *Thermincola* and e) *abcA*.

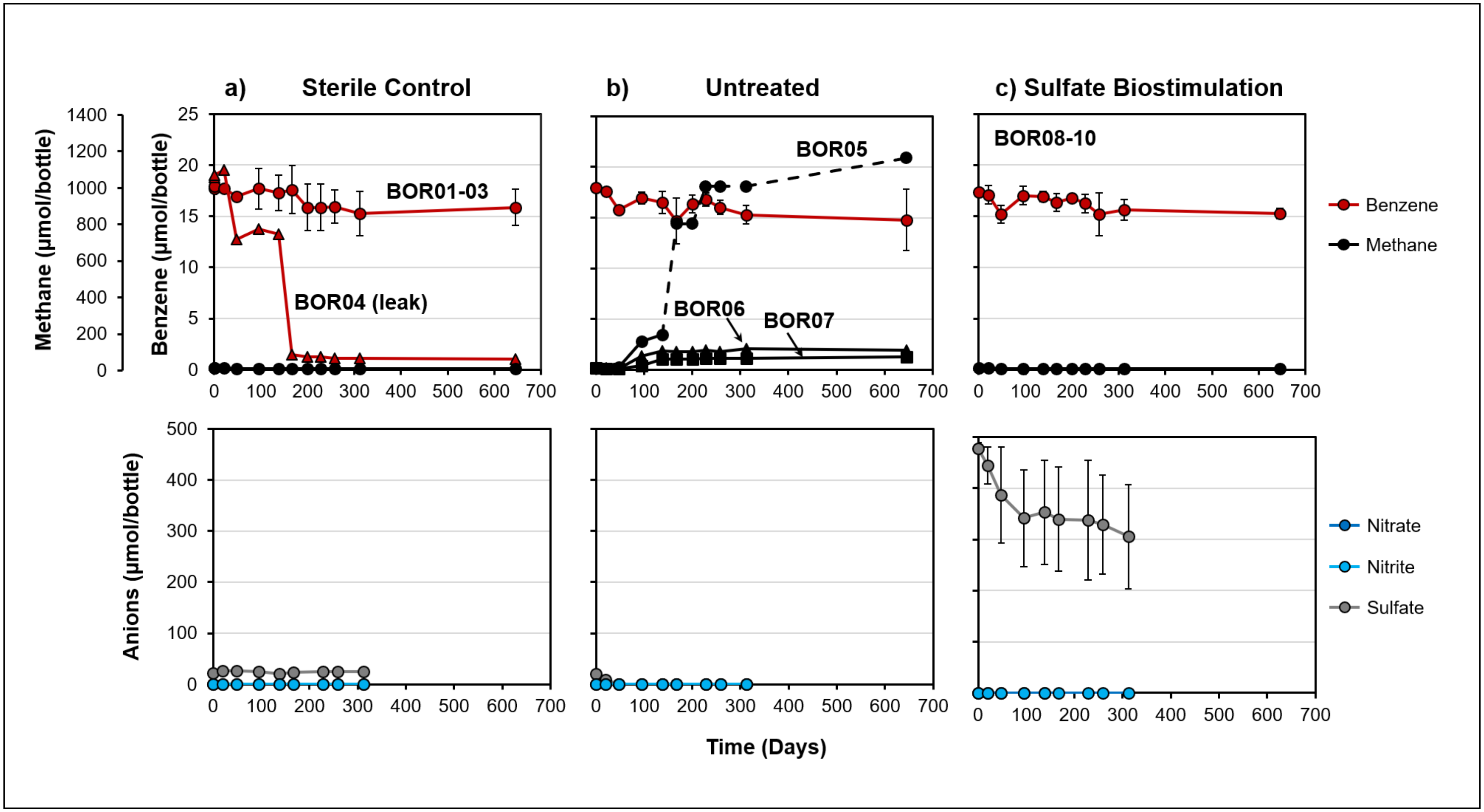


**Figure S2.** Benzene (top panels) and anion (bottom panels) degradation profiles of a) mercuric chloride-poisoned sterile control bottles and active microcosms that were b) left untreated or c) amended with 2 mM sulfate. Most results shown are the average of 3 or 4 replicates (error bars = standard deviation) over 645 days of incubation; replicates with more variable results are plotted separately. Benzene losses in BOR04 were abiotic and due to a leaky cap. The cap was replaced on Day 167 and no additional benzene loss was observed. Dashed lines represent methane datapoints that exceeded liquid saturation limits and reported values may be inaccurate. Anion data was not collected on Day 645.

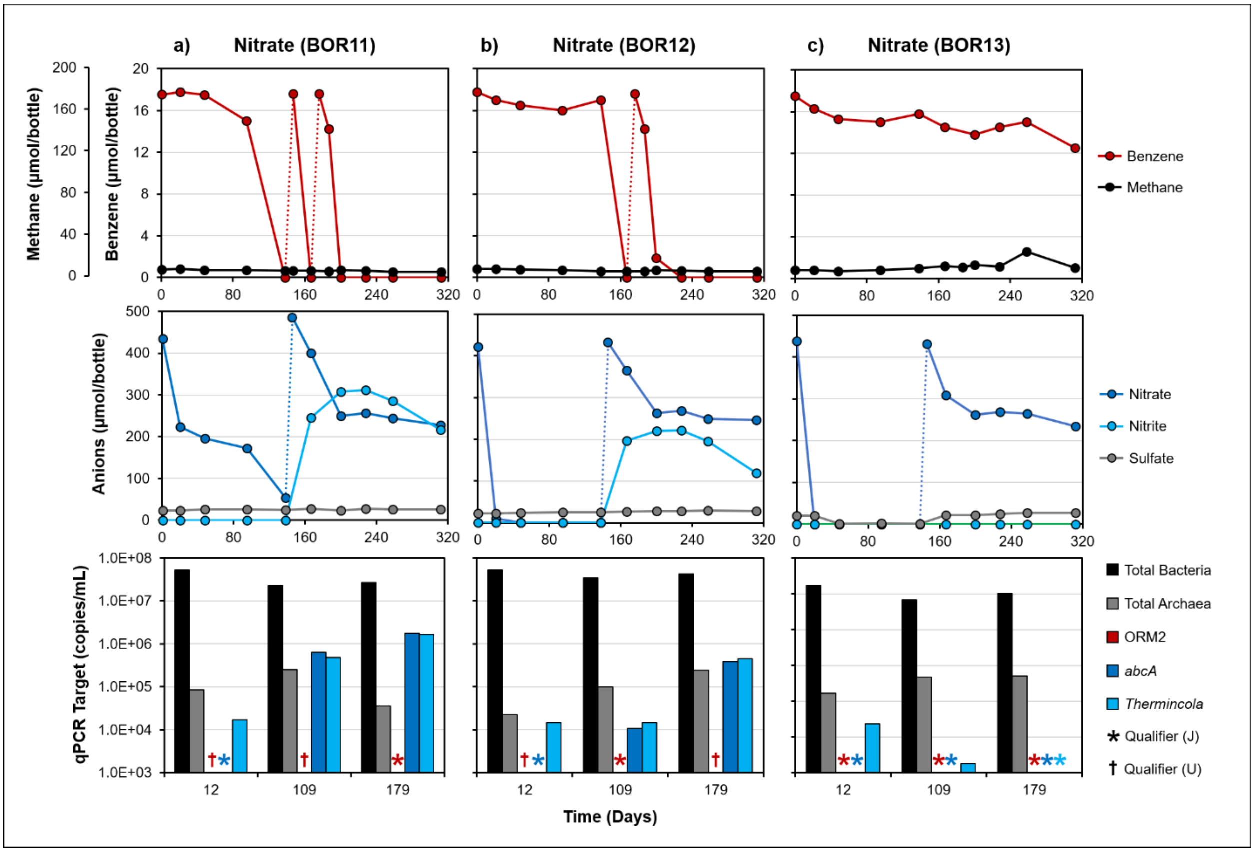


**Figure S3.** Benzene (top panels) and anion (center panels) degradation profiles of active bottles amended with 2 mM nitrate. Electron donor and electron acceptor refeeding events are marked with dotted lines. The bottom panels summarize the abundances of targeted 16S rRNA gene copies and *abcA* for each microcosm. qPCR targets below quantifiable limits (< $10^3$ copies/mL) or below detection are designated by J and U qualifiers, respectively. Each replicate is shown individually.

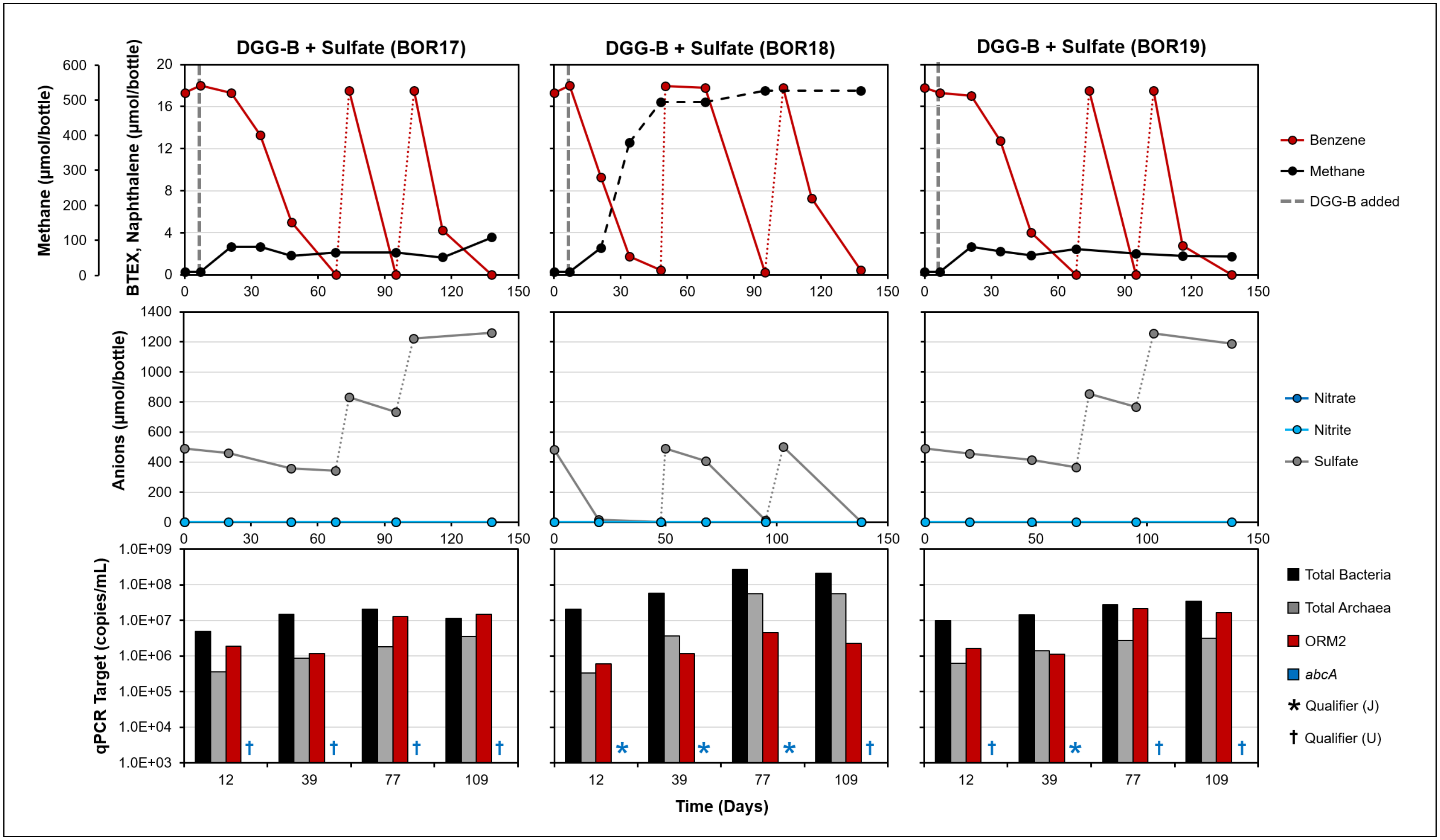


**Figure S4.** Benzene (top panels) and anion (center panels) degradation profiles of active amended with 2.5% *v/v* DGG-B and 2 mM sulfate. Electron donor and electron acceptor refeeding events are marked with dotted lines. Dashed lines represent methane datapoints that exceeded liquid saturation limits and reported values may be inaccurate. The bottom panels summarize the abundances of targeted 16S rRNA gene copies and *abcA* for each microcosm. qPCR targets below quantifiable limits (< $10^3$ copies/mL) or were undetectable are designated by J and U qualifiers, respectively. Data for each replicate is shown individually.

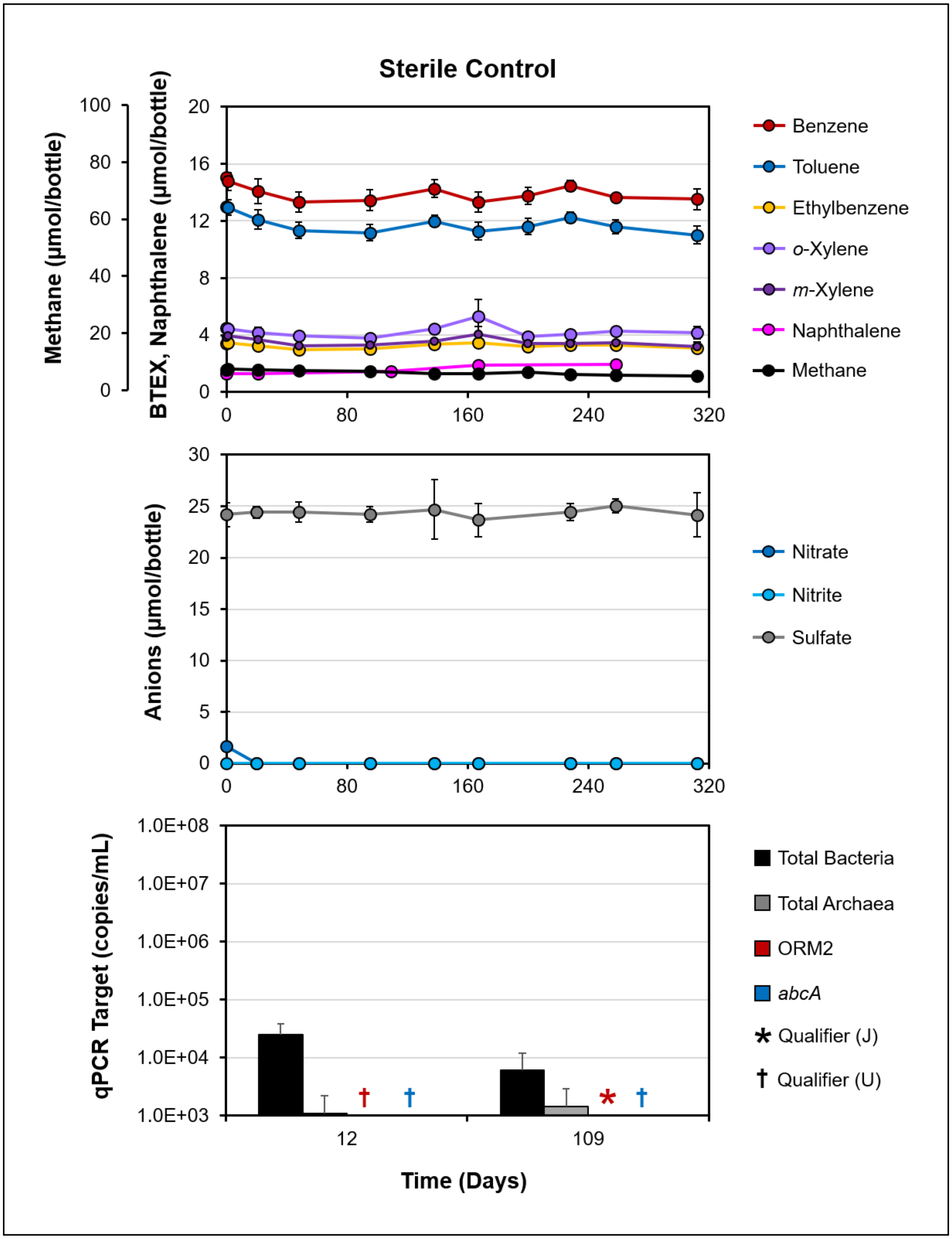


**Figure S5.** Mixed hydrocarbon (top panels) and anion (center panels) degradation profiles in bottles poisoned with mercuric chloride. The bottom panels summarize the abundances of targeted 16S rRNA gene copies and *abcA* at all available molecular timepoints. Results shown are the average of four replicates (error bars = standard deviation). qPCR targets below quantifiable limits (< $10^3$ copies/mL) or below detection are designated by J and U qualifiers, respectively. Naphthalene data was collected on Day 435 (not shown) and no changes were observed.

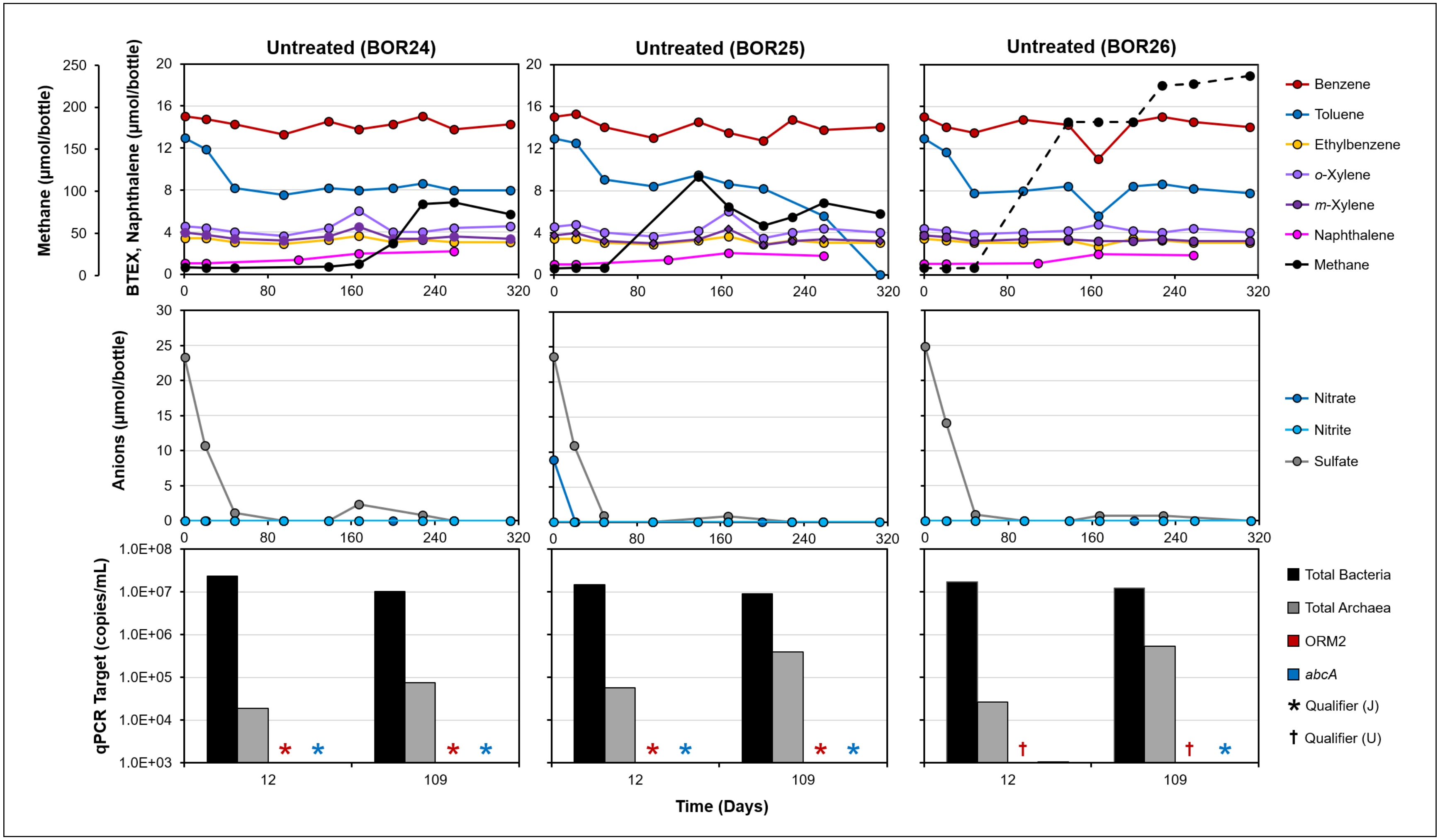


**Figure S6.** Hydrocarbon (top panels) and anion (center panels) degradation profiles of active bottles that were left untreated, simulating natural attenuation. Data for each replicate is shown individually. Dashed lines represent methane datapoints that exceeded liquid saturation limits and reported values may be inaccurate. The bottom panels summarize the abundances of targeted 16S rRNA gene copies and *abcA* for each microcosm. qPCR targets below quantifiable limits (< $10^3$ copies/mL) or were below detectable limits are designated by J and U qualifiers, respectively. Naphthalene data was collected on Day 435 (not shown) and no changes were observed.

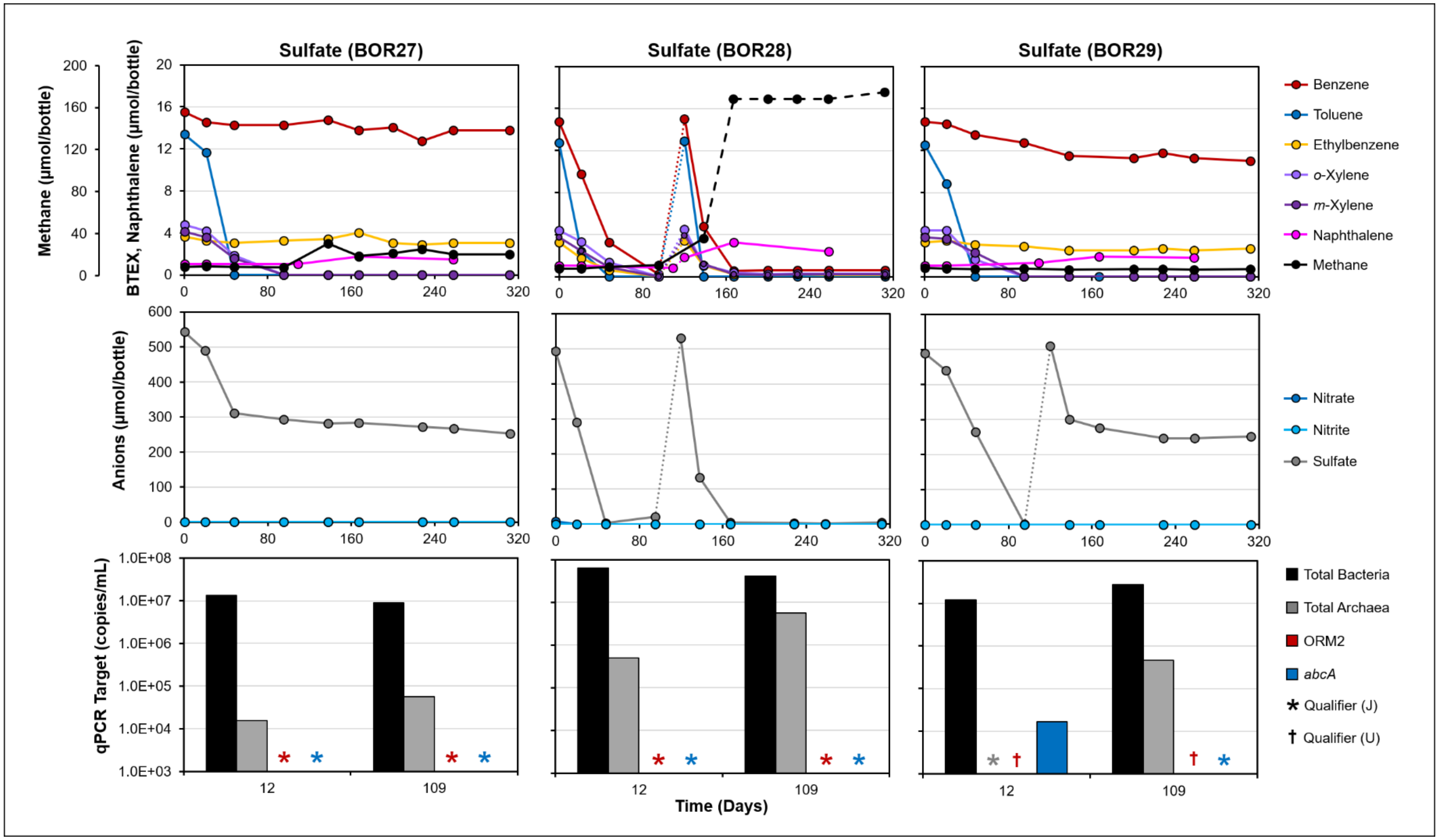


**Figure S7**. Hydrocarbon (top panels) and anion (center panels) degradation profiles of active bottles amended with 2 mM sulfate. Electron donor and electron acceptor refeeding events are marked with dotted lines. Dashed lines represent methane datapoints that exceeded liquid saturation limits and reported values may be inaccurate. Some BTEX losses in BOR28 were likely abiotic and attributed to a leaky cap, replaced on Day 167. The leaky cap may also allowed for hydrogen ingress from the glovebox, contributing to higher than expected nitrate consumption and excessive methane production. The bottom panels summarize the abundances of targeted 16S rRNA gene copies and *abcA* for each microcosm. qPCR targets below quantifiable limits (< $10^3$ copies/mL) or below detectable limits are designated by J and U qualifiers, respectively. Naphthalene data was collected on Day 435 (not shown) and no changes were observed.

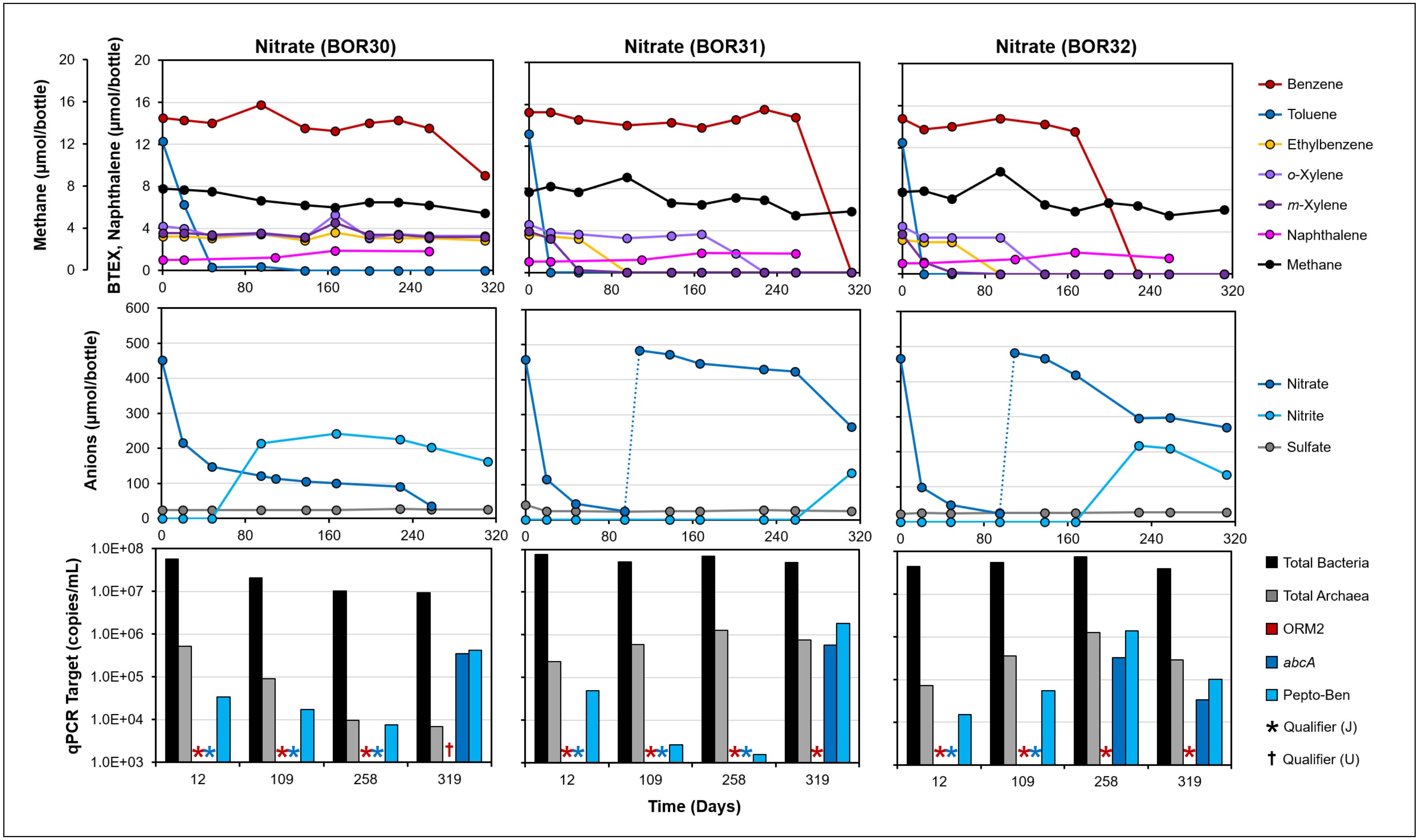


**Figure S8.** Hydrocarbon (top panels) and anion (center panels) degradation profiles of active bottles amended with 2 mM nitrate. Electron acceptor refeeding events are marked with dotted lines. The bottom panels summarize the abundances of targeted 16S rRNA gene copies and *abcA* for each microcosm. qPCR targets below quantifiable limits (< $10^3$ copies/mL) or were undetectable are designated by J and U qualifiers, respectively. Naphthalene data was collected on Day 435 (not shown) and no changes were observed.

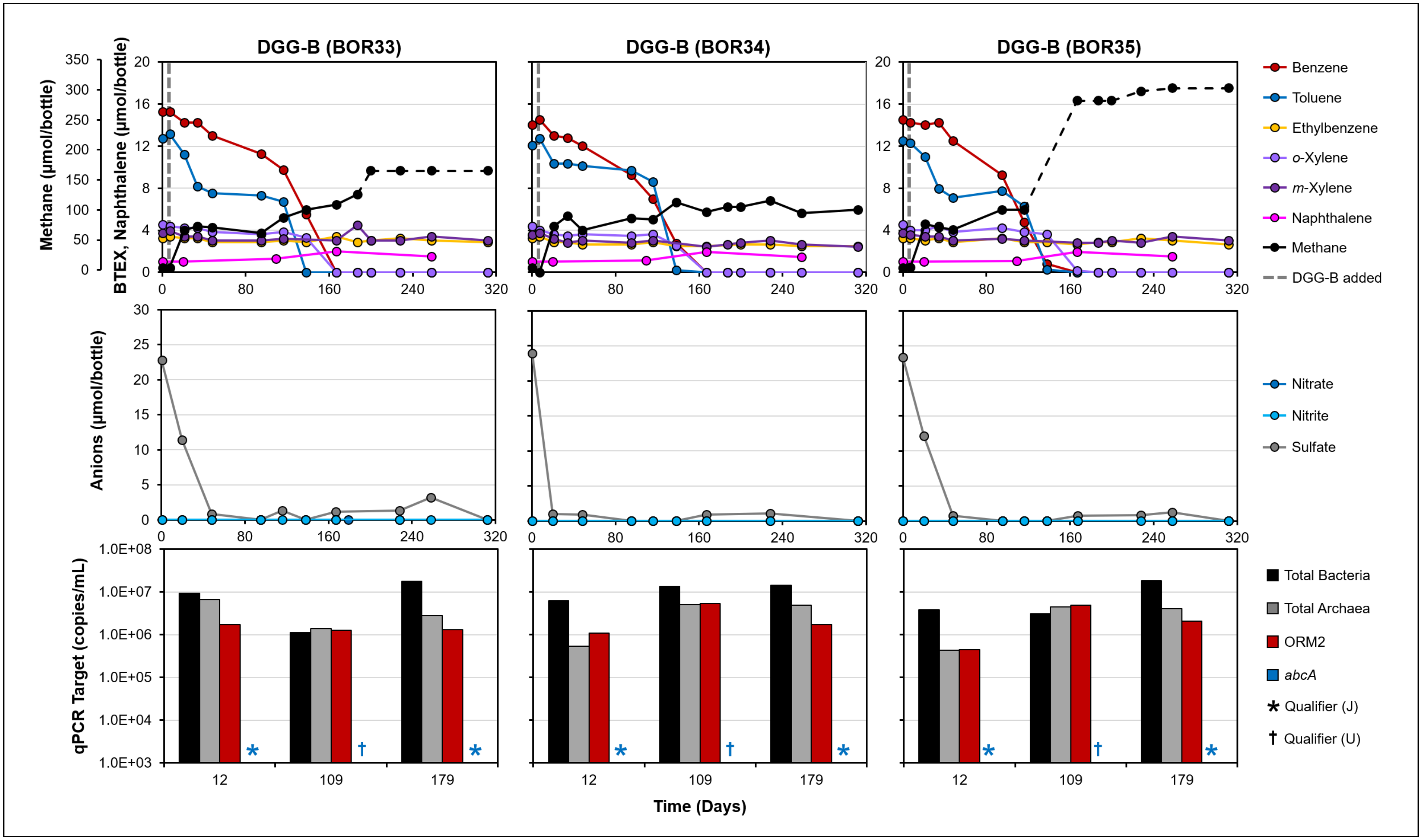


**Figure S9.** Mixed hydrocarbon (top panels) and anion (center panels) degradation profiles of active bottles amended with 2.5% *v/v* DGG-B culture. The bottom panels summarize the abundances of targeted 16S rRNA gene copies and *abcA* for each microcosm. Dashed lines represent methane datapoints that exceeded liquid saturation limits and reported values may be inaccurate. qPCR targets below quantifiable limits (< $10^3$ copies/mL) or were undetectable are designated by J and U qualifiers, respectively. Naphthalene data was collected on Day 435 (not shown) and no changes were observed. Data for each replicate is shown individually.

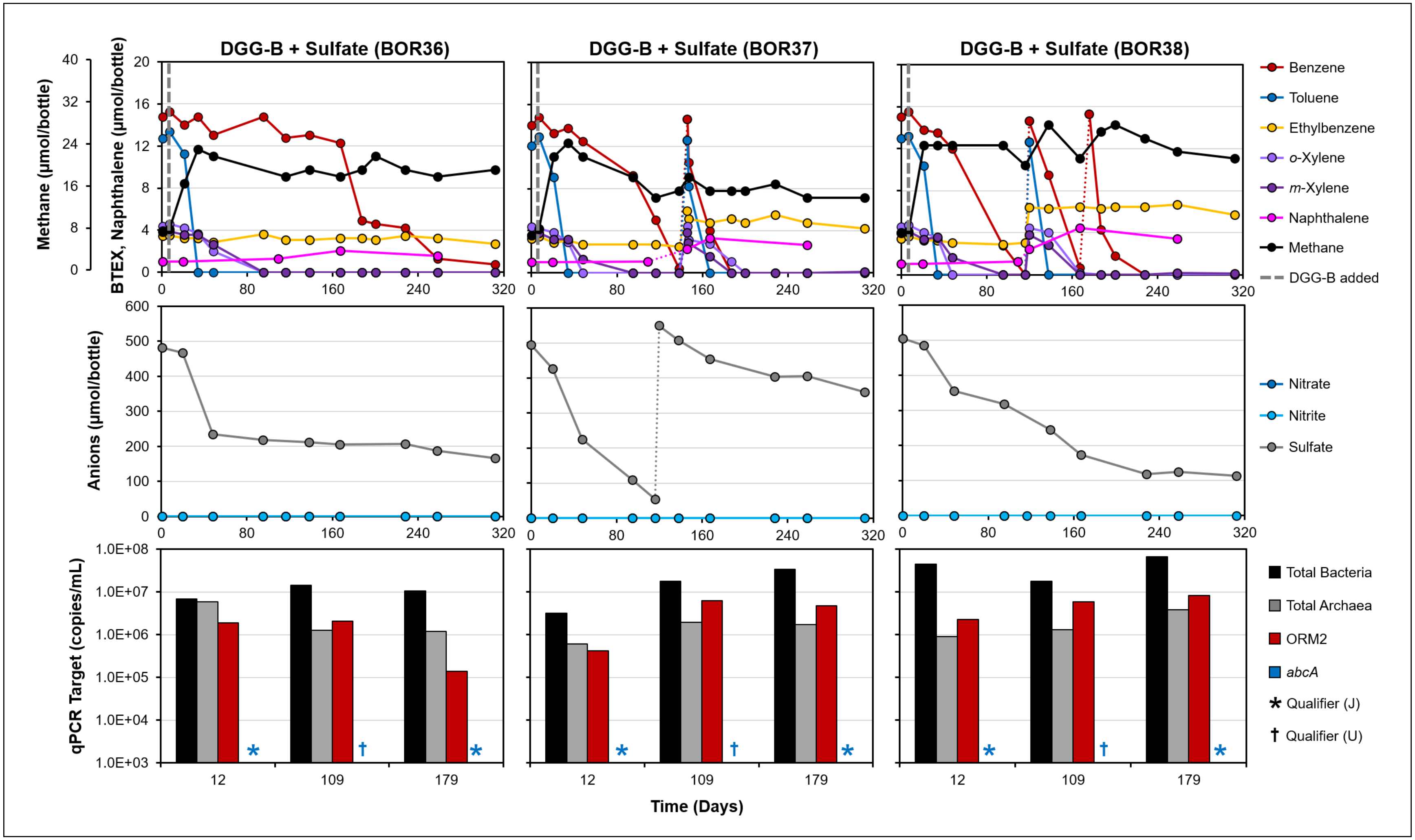


**Figure S10.** Mixed hydrocarbon (top panels) and anion (center panels) degradation profiles of active bottles inoculated with 2.5% *v/v* DGG-B culture and amended with 2 mM sulfate. Electron donor and electron acceptor refeeding events are marked with dotted lines. The bottom panels summarize the abundances of targeted 16S rRNA gene copies and *abcA* for each microcosm. qPCR targets below quantifiable limits (< $10^3$ copies/mL) or were undetectable are designated by J and U qualifiers, respectively. Naphthalene data was collected on Day 435 (not shown) and no changes were observed. Data for each replicate is shown individually.

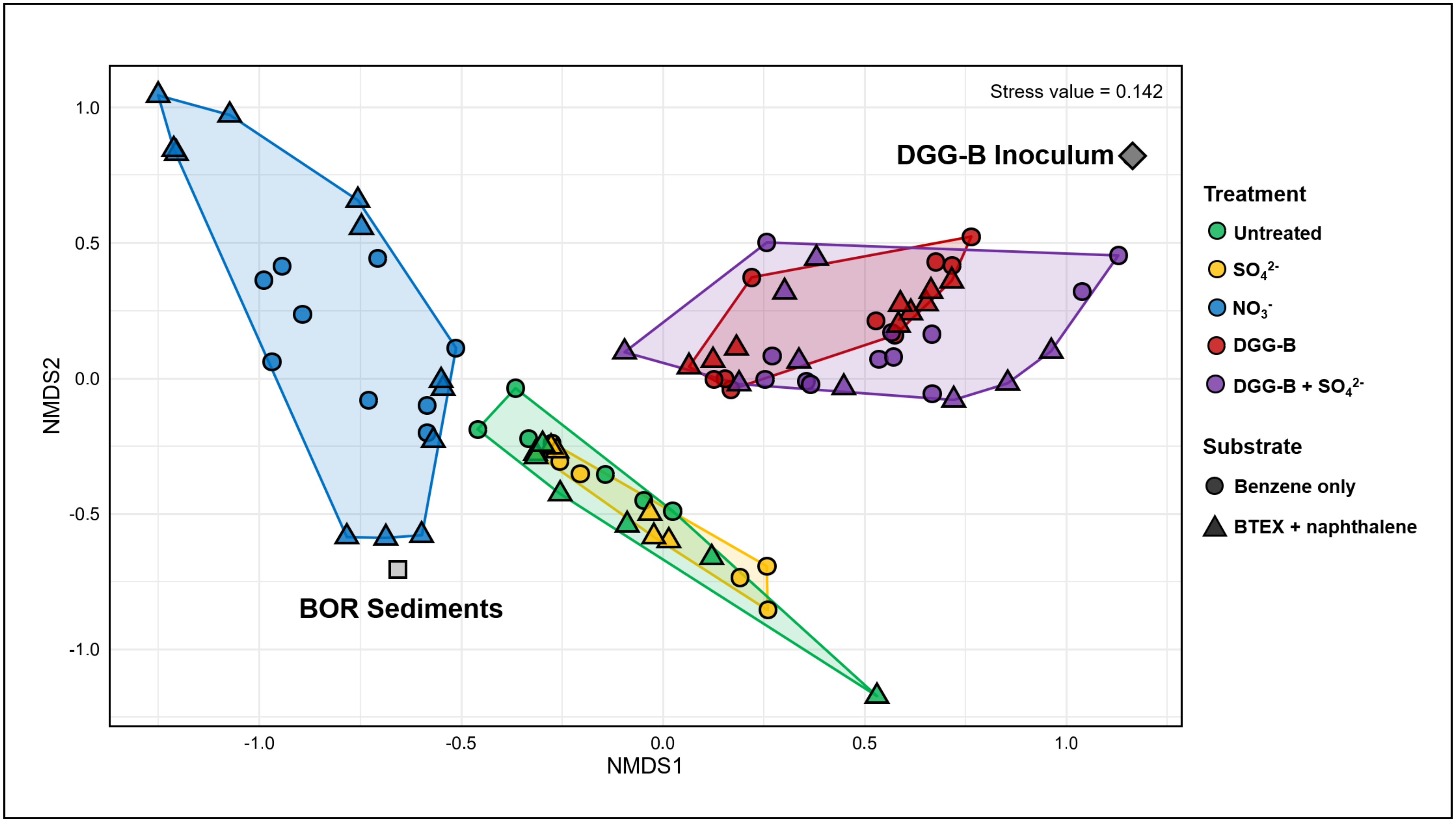


**Figure S11:** Bray-Curtis NMDS plot of all 16S rRNA gene amplicon samples what date. ASVs with an abundance of < 10 reads/sample were removed from the final dataset. A plot with the same overall clustering was obtained when including all ASVs (not shown).

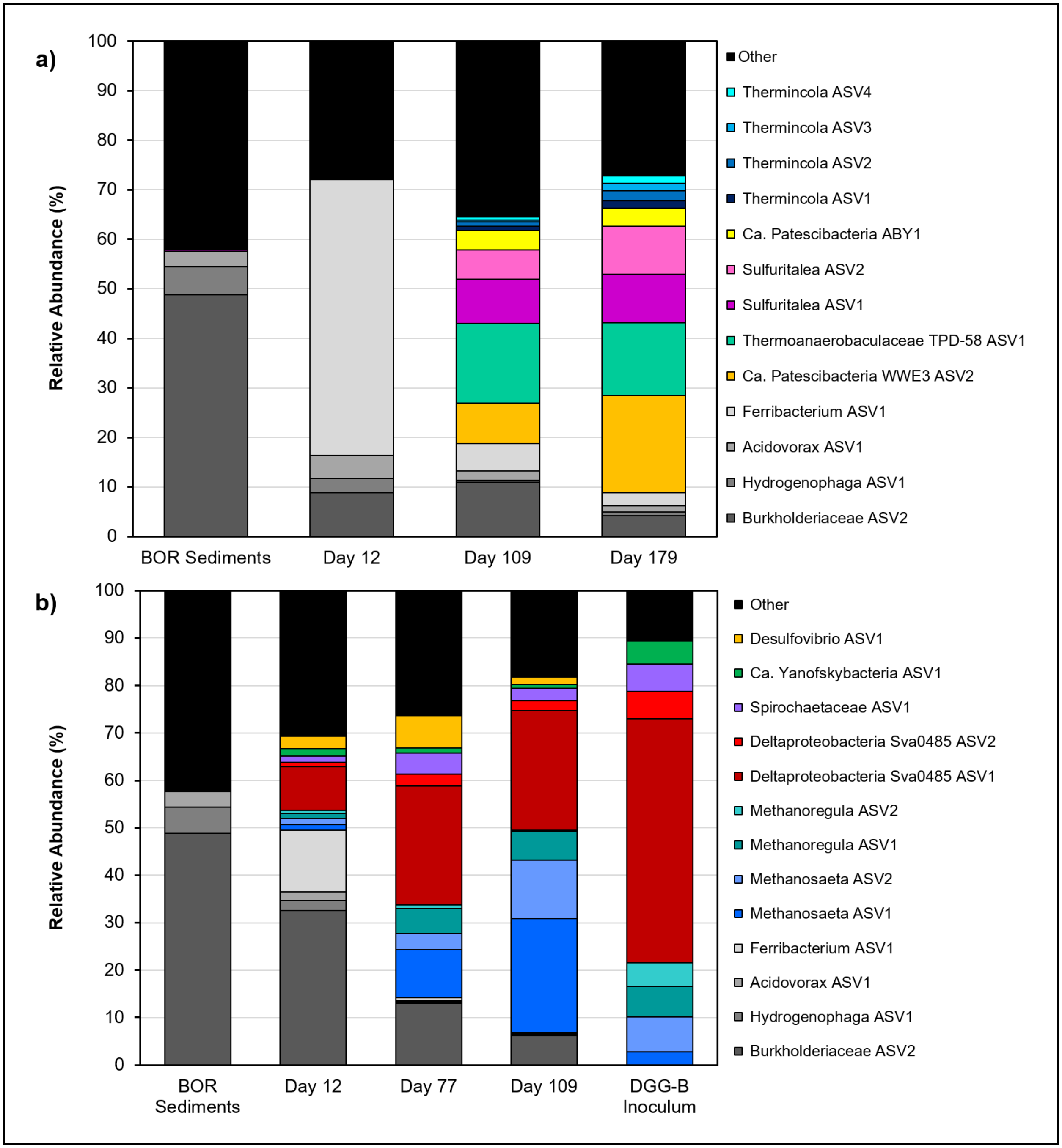


**Figure S12**. Time course microbial community composition of a) nitrate biostimulation microcosm BOR11 and b) DGG-B bioaugmentation microcosm BOR16, relative to original Borden sediments and DGG-B inoculum, respectively.

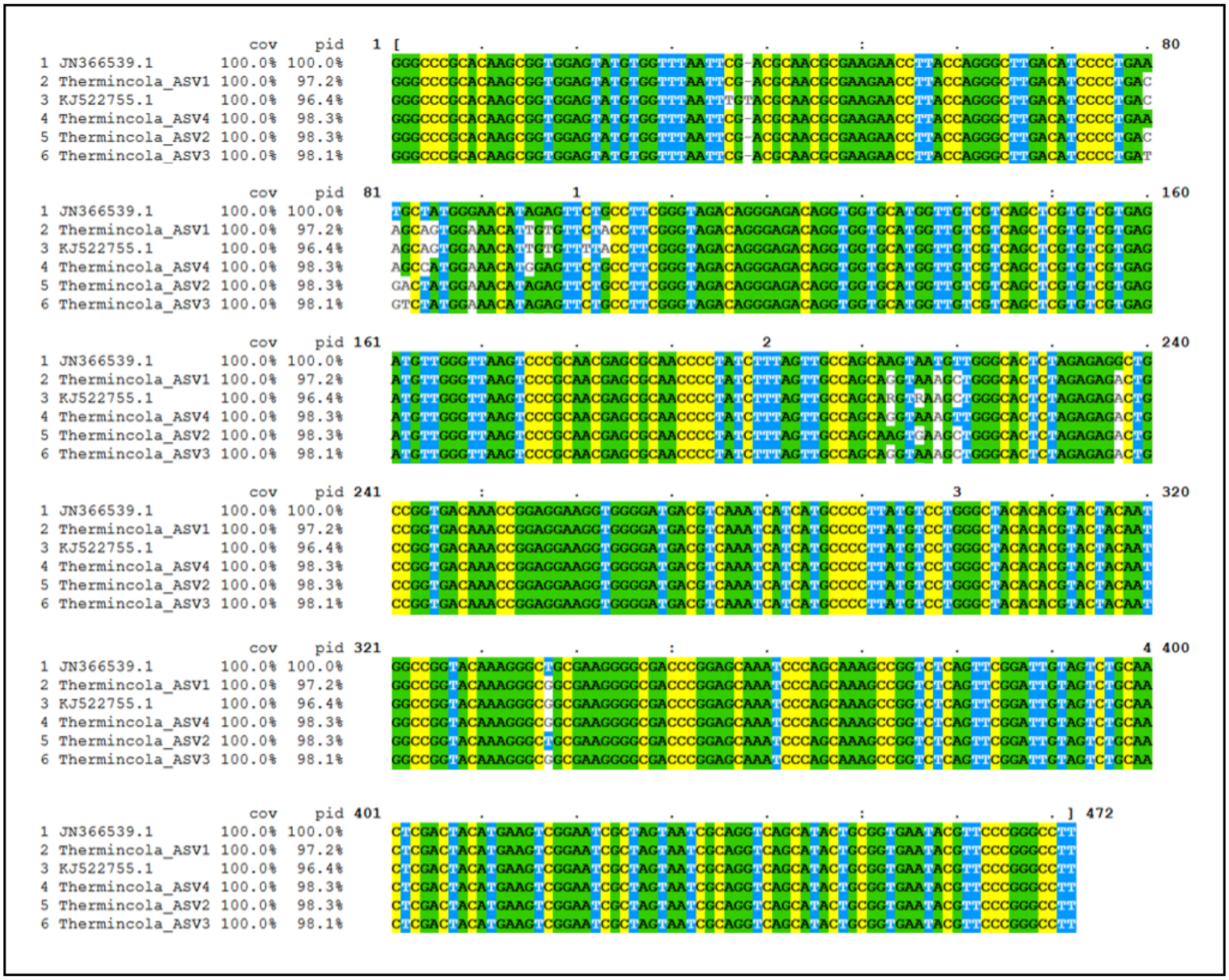


**Figure S13:** Multiple sequence alignment of *Thermincola* ASV1-4 (from BOR sediments) against two reference 16S rRNA gene sequence clones from nitrate-reducing, benzene-degrading *Thermincola* spp., one from the enrichment culture Cartwright-NO3$^-$ (KJ522755.1) and one from a bioreactor enrichment culture (JN366539.1). Sequences were aligned in MUSCLE [7] and visualized in MView [8]. The MUSCLE program selected JN366539.1 as its reference sequence; percent coverage (cov) and percent identity (pid) values are relative to the *Thermincola* ASV2. Nucleotide positions were coloured by identity.

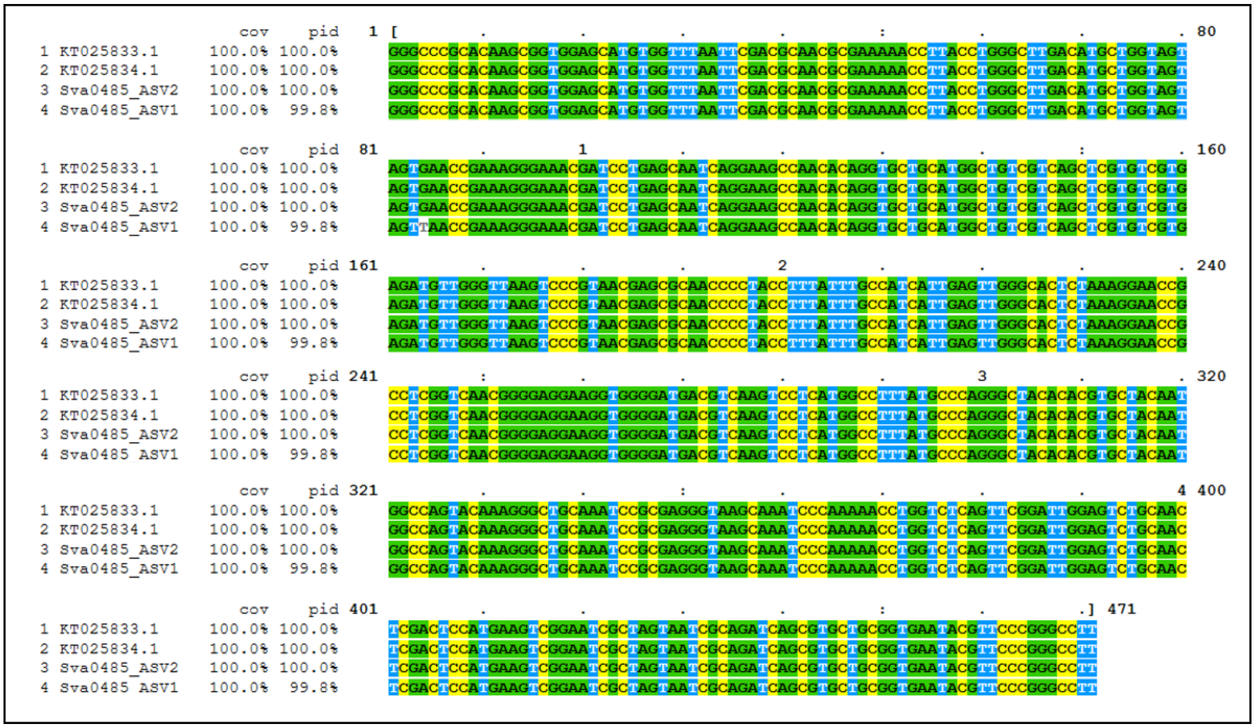


**Figure S14:** Multiple sequence alignment of *Deltaproteobacteria* Sva0485 ASV1-2 against reference 16S rRNA gene sequence clones for *Deltaproteobacteria* ORM2a (KT025833.1) and ORM2b (KT025834.1). Sequences were aligned in MUSCLE [7] and visualized in MView [8]. The MUSCLE program selected KT025833.1 as its reference sequence; percent coverage (cov) and percent identity (pid) values are relative to the Sva0485 ASV2. Nucleotide positions were coloured by identity.
